# Rapid alternation of conscious and suppressed stimulus representations in the primate brain

**DOI:** 10.64898/2026.10.15.759299

**Authors:** Janis K. Hesse, Frank F. Lanfranchi, Conor McGrory, Yuelin Shi, Tatiana A. Engel, Doris Y. Tsao

**Affiliations:** Department of Neuroscience, University of California, Berkeley, Berkeley, CA, USA; Princeton Neuroscience Institute, Princeton University, Princeton, NJ, USA; Division of Biology and Biological Engineering, California Institute of Technology, Pasadena, CA, USA; Howard Hughes Medical Institute, Berkeley, CA, USA

## Abstract

The neural correlates of consciousness remain elusive^1^. Previous studies found that high-level visual cortex is weakly modulated by conscious perception^2,3^, yet a pure signal of conscious perception has not been found anywhere in the brain. Here, we employed Neuropixels recordings across multiple face patches in inferotemporal and prefrontal cortex during a binocular flash suppression paradigm^4^, in which one of two images is consciously perceived while the other, though continuously present, is suppressed from awareness. We found that, during a constant stimulus and percept, the brain rapidly alternates, about 15 times per second, between conscious and suppressed micro-epochs. During conscious micro-epochs, a pure signal of conscious perception was present, allowing accurate reconstruction of the conscious image from neural activity. During suppressed micro-epochs this reversed: the pure signal now tracked the suppressed stimulus, allowing accurate reconstruction of the suppressed image. At the single-cell level, neurons responded selectively to the conscious stimulus and not to the suppressed stimulus during conscious micro-epochs; conversely, during suppressed micro-epochs, neurons responded selectively to the suppressed stimulus. Despite this apparent symmetry, the two epochs could be reliably distinguished, with feature tuning amplified during conscious micro-epochs. Recordings in prefrontal cortex yielded qualitatively similar results. Together, these findings show that both inferotemporal and prefrontal cortex rapidly interleave conscious with suppressed content, indicating that conscious access requires temporal selection beyond localization.

## Introduction

At any moment, only some of the brain’s activity reaches conscious awareness, while the rest is processed without ever being experienced. How this division arises is among the most basic unsolved problems in neuroscience. We set aside the “hard problem” of why neural activity feels like anything, and ask: what distinguishes neural activity that is consciously accessible from activity that is not?

Answering this question requires a situation in which the physical input stays fixed while conscious perception changes, so that differences in neural activity can be attributed to perception rather than to the stimulus. Binocular rivalry is a paradigm that provides exactly this dissociation^5,6^. When conflicting images are presented separately to the two eyes, observers consciously perceive only one image at a time, despite both images remaining physically present. In binocular flash suppression, the dissociation is made experimentally controllable^3,4,7^: when one image is presented alone to one eye, and then a second image is abruptly introduced to the other eye, observers typically perceive the new image (**Fig. 1a**).

**Figure 1.**
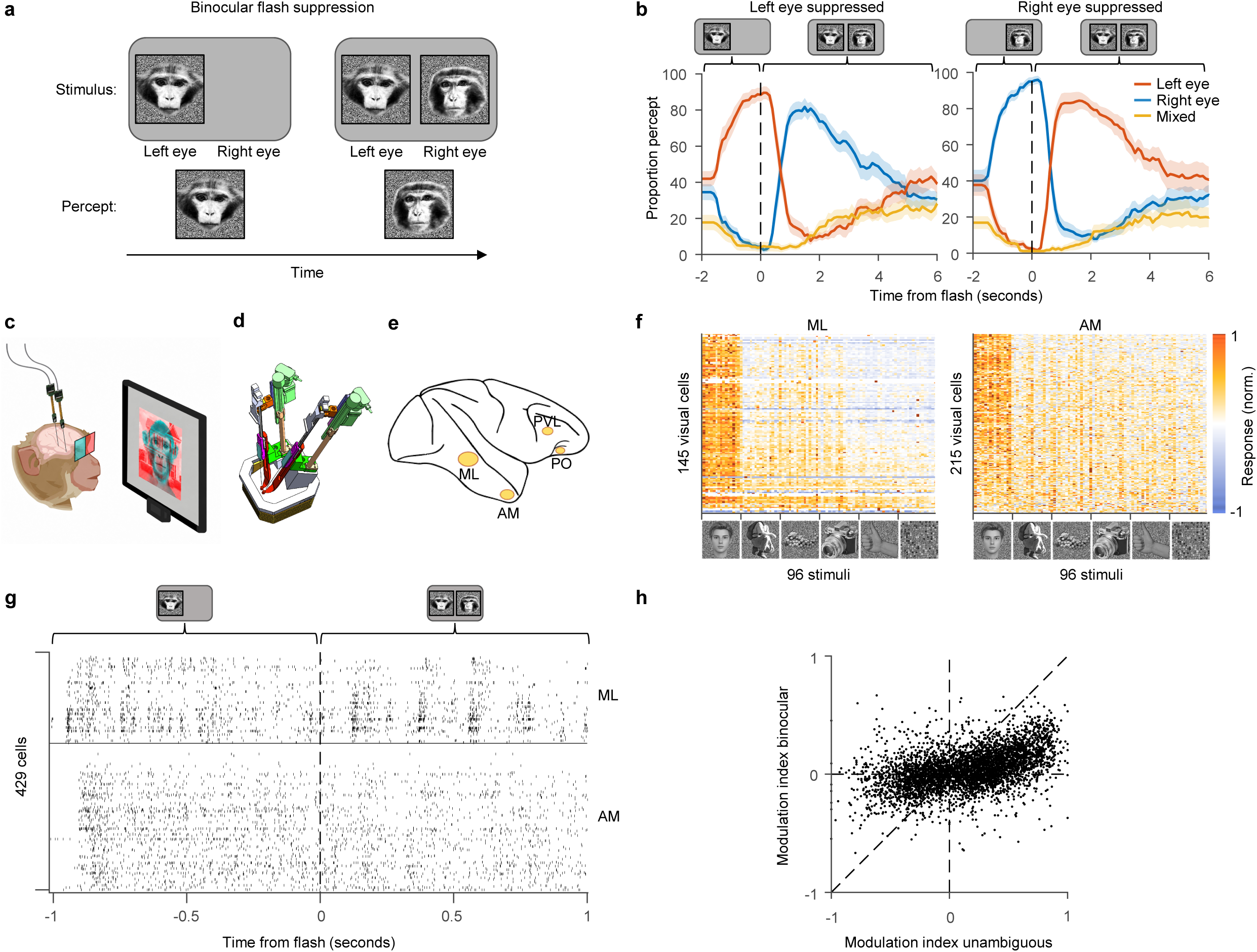
Large-scale recordings across cortical regions during a binocular flash suppression paradigm. [Note: All human faces in this figure have been replaced by synthetically generated faces due to bioRxiv policy on displaying human faces; they are not photographs.] (**a**) Schematic of binocular flash suppression paradigm. First, a priming stimulus is presented monocularly in one eye. Then, without a blank period, a different stimulus is flashed to the other eye, and both are presented binocularly. Following the flash, the flashed stimulus is consciously perceived, while the priming stimulus is perceptually suppressed. (**b**) Perceptual reports in human participants verify perceptual suppression. Human participants were instructed to report whether they were seeing the stimulus in the left eye, right eye or mixed. The priming stimulus was presented at t=0 seconds in the left eye (left) and right eye (right), respectively. The dominant stimulus was flashed in the other eye at 2 seconds, causing the conscious percept to flip on most trials. Curves indicate average proportion of reported percepts of the stimulus in the left eye (red), right eye (blue), or a mixed percept (yellow). We compared the difference in report probability between the flashed and the primed stimulus, averaged over left eye and right eye primed trials, 300–1000 ms + report latency after flash onset with a two-tailed t-test against 0 (t(13) = 9.4, p = 4 × 10^−7^, n = 14 participants, see Methods). (**c**) Schematic of recording setup. Monkeys viewed flash suppression stimuli on a screen through anaglyph goggles to separate input to the left and right eyes while two Neuropixels probes were inserted into different nodes of the visual cortical hierarchy. (**d**) CAD drawing of Neuropixels setup. Neuropixels probes were advanced using a custom insertion mechanism designed to target the desired regions. (**e**) Schematic of macaque brain with four regions targeted for electrophysiological recordings. (**f**) Response profiles of visually responsive cells (rows) to 96 visual stimuli (columns) from six categories, including faces and other objects. Shown are example sessions from ML (left), and AM (right). (**g**) Raster plot showing an example flash suppression trial. Black ticks represent spikes of 429 simultaneously recorded cells (y-axis) from ML and AM. Vertical bars and icons show when the priming stimulus and the rivalry stimulus were presented, respectively. (**h**) Scatter plot showing modulation indices for the binocular (y-axis) versus unambiguous (x-axis) conditions. Each dot corresponds to a cell and stimulus pair where the responses were significantly selective for the unambiguous stimulus. Modulation during the unambiguous condition was compared to modulation during the binocular condition using a two-sided paired t-test (*t*(5549) = 25.2, *p* = 8 × 10^−133^, *n* = 5550 cell-stimulus pairs).

Rivalry studies have shown that neural activity in the primate brain is modulated by conscious perception: a cell that responds more to faces than houses will, under rivalry, fire more when the face is perceived than when the house is perceived, even though both images are continuously present. Yet this perceptual modulation poses a persistent puzzle: it has consistently been found to be weaker than the modulation the same cell shows when the face and house are simply presented one at a time^2,3,8,9^. Moreover, neurons continue to carry information about the suppressed image^2^. Thus, previous studies *have not revealed a pure neural representation of conscious perception, i.e., one uncontaminated by suppressed representation and closely resembling the representation of a perceptually-equivalent unambiguous stimulus, in any recorded region of the brain during rivalry*.

This absence of a pure correlate has contributed to an ongoing debate over whether the neural correlates of consciousness reside in the front or the back of the brain, with frontal theories such as Global Neuronal Workspace and Higher-Order theories on one side and posterior theories such as Integrated Information Theory and Recurrent Processing Theory on the other^10^. Recent large-scale efforts, including the COGITATE adversarial collaboration^11^, have tested these competing predictions, yet no definitive anatomical locus has emerged.

Conscious perception is a dynamic process. A given sensory input can be interpreted in multiple ways, and perception actively resolves which interpretation we experience at any moment^12^. Rivalry makes this especially stark: with the physical input held constant, perception settles into one of two interpretations and alternates over time. Understanding this process therefore requires asking *how different stimulus representations are dynamically organized in time* within cortical populations. Previous studies using conventional single-unit recordings have generally required averaging across trials, because the activity of individual neurons is too sparse and variable to resolve moment-to-moment representational dynamics within a single trial. Addressing this requires tracking the coordinated activity of many neurons in real time while presenting enough stimuli to characterize what each momentary population state represents.

Here, we examined how conscious and suppressed representations are organized in time, using large-scale Neuropixels recordings^13^ from macaque face patches in inferotemporal and prefrontal cortex during binocular flash suppression. Presenting a large set of face stimuli allowed us to reconstruct the specific face stimuli represented by the population^14^, while the use of Neuropixels probes allowed us to track changes in representations within individual trials, at a resolution not accessible with single-neuron recordings. We found that even during a stable percept and constant stimulus, inferotemporal population activity rapidly alternated between two states, one purely representing the consciously perceived face and the other purely representing the suppressed face. Conscious versus suppressed face epochs could be reliably decoded, with higher gain for representation of the conscious face during conscious epochs than for the suppressed face during suppressed epochs. Prefrontal cortex showed the same interleaved structure. These findings show that conscious and suppressed representations are temporally multiplexed within cortical population activity; both inferotemporal and prefrontal cortex carry a rich representation of suppressed stimuli that are not consciously perceived, indicating that selective access to the conscious percept requires a temporal selection mechanism.

## Results

### Binocular flash suppression dissociates conscious perception from the physical stimulus

To dissociate conscious perception from the physical stimulus, we used binocular flash suppression (**Fig. 1a**). A priming image is shown to one eye, and after an extended period (1 or 2 s) a second image is flashed to the other eye; the flashed image dominates perception while the priming image is suppressed from awareness.

We first verified the effectiveness of this paradigm in human participants. Participants reported by button press whether they perceived the image presented to the left eye, the image presented to the right eye, or a mixed percept. Shortly after the flash, participants predominantly reported perceiving the newly introduced stimulus, with few reports of the priming or mixed percept (*p* = 4 × 10^−7^, t-test on proportion of flashed versus priming percept across 14 participants, **Fig. 1b**, see Methods). For all subsequent analyses, we focus on the period 300 to 1000 ms after the flash, where the flashed stimulus dominated perception and the priming stimulus was suppressed.

### Average neural activity is only weakly modulated by conscious perception

We performed large-scale electrophysiological recordings while rhesus macaque monkeys viewed the same flash suppression displays verified in humans (**Fig. 1c**). NHP Neuropixels probes were inserted into multiple nodes of the macaque brain using a custom insertion mechanism (**Fig. 1d**): face patches ML and AM in inferotemporal cortex, and the orbitofrontal (PO) and ventrolateral (PVL) regions of prefrontal cortex. Data were recorded from two monkeys; throughout the paper, we show data pooled across the two animals in the main figures; data for individual animals are shown in the extended data. Face patches were localized with fMRI using a faces-versus-objects contrast (**Extended Data Fig. 1**). A majority of cells in ML and AM were face-selective (**Fig. 1f**, 91% and 71% of visually responsive cells for the example session shown; faces vs. objects, t-test, *p* < 0.05). A single trial of simultaneously recorded ML and AM activity is shown in **Fig. 1g**.

We first focus on inferotemporal cortex. We asked whether single neurons there purely represented the consciously perceived stimulus. For each cell, we compared its responses to the two images of a stimulus pair, A and B, using a modulation index 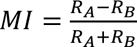, where *R* and *R* are the cell’s average firing rates to stimulus A and stimulus B. We compared modulation during rivalry with modulation during an unambiguous alternation condition. During the unambiguous condition, stimuli were presented alone after priming rather than in binocular conflict, providing a measure of the cell’s visual selectivity. During binocular flash suppression the same two stimuli were presented in conflict, and we computed the index with respect to the consciously dominant stimulus, so that *R_A_* and *R_B_* now reflect responses when stimulus A or stimulus B, respectively, was perceived. If activity during flash suppression purely reflected the conscious percept, modulation in the binocular condition would match modulation during unambiguous priming. Instead, modulation during binocular flash suppression was much weaker than during unambiguous alternation (**Fig. 1h**, *p* = 8 × 10^−133^, paired t-test over 5550 cell-stimulus pairs). Virtually all neurons that strongly discriminated the two stimuli during unambiguous presentation showed much smaller differences when the same stimuli were presented in binocular conflict, with nearly all points falling below the identity diagonal (slope = 0.23). This weak modulation is puzzling: perceptually, one image is clearly seen while the other is absent from awareness, yet the neural responses only weakly distinguished the two.

A clue to the origin of this weak modulation comes from an earlier finding that the suppressed stimulus, though absent from awareness, can still be decoded from inferotemporal activity during rivalry^2^. In that study, however, only three stimuli were used (and previous studies generally contrasted only two rival alternatives^3,8,9,15^), leaving opaque what, if any, details of either the conscious or the suppressed face are represented during rivalry. We therefore set out to characterize the detailed content of conscious and suppressed representations. The face patch system is well suited to this: face identity can be described in a parametric feature space to which cells are linearly tuned, so that face features can be reconstructed from neural responses by linear regression^14^. To precisely map the feature code of each cell, we presented 1500 faces in flash suppression. This allowed us to ask not merely whether a conscious or suppressed face was decodable, but precisely which features of each were represented at each moment.

### Single-trial population activity can be partitioned into rapidly alternating states

Having access to the represented content at each moment in time, we first asked whether there are epochs in which only the conscious or only the suppressed face is represented. First, we computed template vectors for the conscious and suppressed faces by taking the average neural responses during priming trials when each was shown alone. For each 10 ms time bin during rivalry, we then asked which template the observed spike counts more probably arose from (a Poisson log-likelihood ratio (LLR) decoder, see Methods), labelling the bin state 1 when the conscious face template was more likely and state 2 when the suppressed face template was more likely (**Fig. 2a**). This produced a time-resolved partition of each trial into two states. During priming, when only one face was present, activity was assigned mostly to the state corresponding to the priming image (**Fig. 2b**). After the flash, however, the population rapidly alternated between the two states, and state 1 and state 2 were occupied about equally (**Fig. 2c**). To quantify how quickly the population alternated, we measured the autocorrelation of the moment-by-moment log-likelihood-ratio signal (**Fig. 2d**, black). It decayed with a time constant of 67 ms, whereas the same signal computed after shuffling time points decayed within a single bin (**Fig. 2d**, grey). Thus, the state transitions occurred more than an order of magnitude faster than the perceptual switches reported during binocular rivalry (see Methods and Results section “Dynamics of state switching” below for more details).

**Figure 2.**
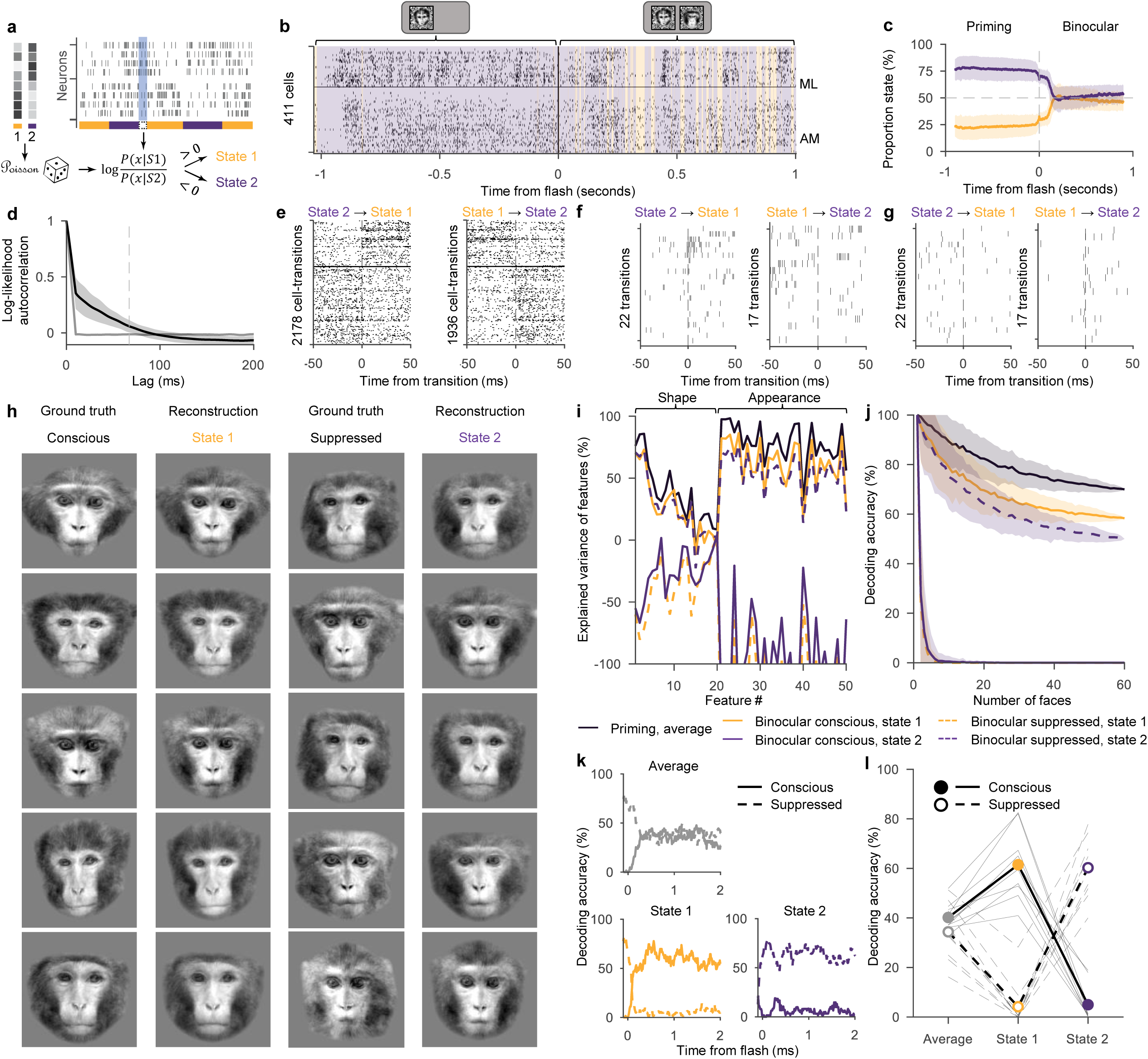
Partitioning neural activity into conscious and suppressed states. (**a**) Schematic of log-likelihood ratio (LLR) state inference method. The LLR finds a partition of the time course into two states. (**b**) Example raster plot and state partition of a trial consisting of a priming stimulus, followed by binocular flash suppression. Orange shading indicates state 1, which corresponds to the flashed, conscious stimulus; purple shading indicates state 2, which corresponds to the suppressed priming stimulus. Vertical line indicates the time of the flash. Horizontal line demarcates 157 selective cells recorded from ML and 254 selective cells recorded from AM. (**c**) Proportion of time points inferred to be state 1 (orange) and state 2 (purple) during priming and binocular condition. Shaded regions indicate standard deviation across sessions. (**d**) Auto-correlation of the log-likelihood ratio for real data (black) and time-shuffled data (grey). Shaded regions indicate standard deviation across sessions. (**e**) Raster plot of neural activity of neurons from ML and AM, aligned to 22 transitions to state 1 (left) and 17 transitions to state 2 (right) during the binocular condition, for an example stimulus pair from the same session as (b). Cells are sorted by their modulation index of the priming responses to the two stimuli. Only cells that significantly preferred one of the stimuli were included. Cells above the solid horizontal line prefer the conscious face, and cells below prefer the suppressed face. Only state transitions where at least 20 ms of one state were immediately followed by at least 20 ms of the other were included. (**f**) Spike raster of an example neuron from ML that preferred the conscious face, from the same session and pair as (e). (**g**) Same as (f) for a simultaneously recorded example neuron from AM that preferred the suppressed face. (**h**) Example reconstructions of face images from neural activity during state 1 and state 2. The first and third columns show the ground truth conscious and suppressed faces presented to the monkey. The second and fourth columns show reconstructions obtained through linear regression of face features from neural activity during priming, and applied to neural activity in the binocular condition during state 1 and state 2, respectively. (**i**) Explained variance of conscious and suppressed face features. Explained variance (y-axis) was computed for the 20 principal shape features and 30 principal appearance features by comparing ground truth features to features predicted by linear regression. Orange solid line: Conscious stimulus features from state 1. Orange dashed line: Suppressed stimulus features from state 1. Purple solid line: Conscious stimulus features from state 2. Purple dashed line: Suppressed stimulus features from state 2. Black solid line: Priming stimulus features predicted from average activity during priming. For each stimulus pair, we compared how much closer the reconstruction was to the conscious than the suppressed face between state 1 and state 2, and tested this difference against zero with a paired t-test (*t*(29) = 48.5, *p* = 3 × 10^−29^, *n* = 30 stimulus pairs, see Methods). (**j**) Decoding accuracy of conscious and suppressed faces during state 1 (orange) and state 2 (purple). Euclidean distance was computed between predicted face features and ground truth face features of the conscious/suppressed face and a given number of randomly selected distractor faces (x-axis). A predicted face was defined as decoded correctly when the distance was smaller to the conscious (solid) or suppressed (dashed) face compared to the distractors. The double dissociation of state-resolved decoding accuracy was tested per rivalry pair with a two-sided sign-flip permutation (*p* = 10^−5^, *n* = 30 stimulus pairs). (**k**) Single-trial decoding for an example session. A linear multi-class decoder was trained on priming responses and applied to binocular rivalry responses to discriminate between 10 stimulus identities that were shown in a separate set of experiments. During binocular rivalry, within a 200-ms sliding window on each trial, the decoder used average activity across all time points in the window (top left), or across only the time points in the window assigned to state 1 (bottom left) or state 2 (bottom right), and the decoded identity was compared to the ground truth conscious (solid) or suppressed (dashed) stimulus. (**l**) Single-trial decoding summary across sessions. Activity was averaged across the entire binocular trial (grey), state 1 (orange), or state 2 (purple), and the decoded identity was compared to the ground truth conscious (solid) or suppressed (dashed) stimulus. Thin lines indicate individual sessions. Thick lines indicate average across the ten sessions. Chance performance is 10%. The difference in decoding accuracy for conscious versus suppressed was evaluated using a two-sided paired t-test (state 1: *t*(9) = 11.0, *p* = 2 × 10^−6^, *n* = 10 sessions; state 2: *t*(9) = −13.7, *p* = 3 × 10^−7^, *n* = 10 sessions).

For stimulus pairs where each of the two faces was preferred by a distinct set of neurons, the alternation was visible both in the population (**Fig. 2e**) and in single-cell rasters (**Fig. 2f, g**).

### Reconstructions from inferred states recover conscious and suppressed stimuli

After partitioning the population activity into epochs whose representations more closely resemble either the conscious or the suppressed stimulus, we next asked what face features were represented during each inferred state. Exploiting the linear tuning of face cells to features of a parametric face space^14^, we trained regression weights to predict face features from responses to priming stimuli (i.e., in the absence of binocular conflict) and then applied these weights to activity during state 1 and state 2 to attempt to reconstruct the face represented in each. We inferred states individually for each of 9 sessions (from two monkeys) and averaged responses of each neuron over all state 1 and, separately, over all state 2 epochs from all trials. We then pooled these state-averaged neural responses from ML and AM neurons across sessions to predict features. The predicted features were transformed into images for visualization.

Strikingly, reconstructions computed from activity during state 1 closely resembled the conscious face, whereas reconstructions computed from state 2 closely resembled the suppressed face (**Fig. 2h**). Thus, even though the suppressed face was not consciously perceived, a clear representation of it was present in IT cells during distinct epochs in time. These example reconstructions suggest that state 1 and state 2 carry separate representations of the conscious and suppressed face. By contrast, a time-averaged null model, in which features were predicted from activity averaged across the binocular period without state segmentation, produced reconstructions that matched neither the conscious nor the suppressed face, but rather their average (**Extended Data Fig. 2**).

To quantify reconstruction accuracy, we computed explained variance of each face feature, defined as the percentage of variance of the true values of that feature across faces explained by its reconstructed values (**Fig. 2i**). During state 1, prediction of the consciously perceived face reached over 80% explained variance for some features. This was only slightly lower than the upper bound for this analysis, obtained by reconstructing faces from average priming responses, which reached over 90% explained variance. The upper bound is informative because priming uses the same regression weights but without binocular conflict, so it shows how well faces can be reconstructed from this population under ideal conditions; state 1 approaches it despite the competing suppressed face. By contrast, explained variance for the suppressed face during state 1 was extremely low, comparable to chance level when that face was not present (**Fig. 2i**, orange dashed trace; compare **Extended Data Fig. 2b**, black dashed trace). Thus, during state 1 there is strong information about the consciously perceived face and little to none about the suppressed face.

During state 2, the pattern reversed: explained variance for the suppressed stimulus was now high, albeit slightly lower than for the conscious face during state 1 (**Fig. 2i, purple**). Explained variance for the conscious stimulus during state 2 fell to chance level as above. Thus, state 1 and state 2 separately encoded the conscious and suppressed faces (*p* = 3 × 10^−29^, t-test, see Methods).

As a complementary measure, we asked whether each reconstruction was accurate enough to identify the presented face among a number of randomly selected faces, following the decoding approach used previously to read out facial identity from face cell responses^14^. For each reconstructed face, we compared its predicted features to the features of the correct face and to those of randomly chosen distractor faces, and counted the reconstruction as correctly decoded when it was closest to the correct face **(Fig. 2j)**. During state 1, decoding accuracy was high for the conscious face and low for the suppressed face; during state 2, this reversed, with high accuracy for the suppressed face and low accuracy for the conscious face (*p* = 10^−5^, permutation test, see Methods).

We also asked whether the state-resolved approach could help with decoding of stimulus identity on *single trials* rather than state-resolved activity averaged across trials in a separate set of 10 experiments where we presented 10 stimuli per experiment. We chose the 10 stimuli from a screening set of 49 stimuli that maximized decodability based on a screening session that was conducted and analysed right before each main session. We presented 25 paired combinations of the 10 stimuli in binocular flash suppression. We then trained a 10-class linear decoder on time-averaged responses on individual priming trials to discriminate the 10 identities. We applied this decoder to single-trial binocular responses: within a 200-ms sliding window on each trial, we decoded separately from the time points assigned to state 1, to state 2, or from both states combined, and asked whether the predicted identity matched the conscious or suppressed stimulus (**Fig. 2k**; see Methods). Across sessions, decoding using average activity during the binocular condition was about equal for the conscious and suppressed stimuli (40% and 34%, respectively). During state 1, on the other hand, the conscious stimulus was decoded far more accurately than the suppressed stimulus (over 61% versus 4%; **Fig. 2l**, *p* = 2 × 10^−6^, paired t-test). During state 2, the pattern reversed (*p* = 3 × 10^−7^, paired t-test).

Together, these results indicate that population activity during rivalry can be partitioned into temporal epochs whose average activity represents either the conscious or the suppressed face. Because the regression weights were trained only on unambiguous priming responses, the accuracy of both reconstructions means that the average activity within each set of epochs reinstates the same feature code evoked when the face is presented alone.

However, because the same neural activity was used both to define the state partition and to evaluate representations within these states, the analyses above are subject to a circular conditioning bias: the circular re-use of the same data samples allows each neuron’s independent spiking fluctuations to influence both state assignment and its subsequent state-conditioned response. The resulting selection bias systematically sorts these fluctuations along the state-decoder direction, artifactually increasing the separation between state-conditioned population averages and inflating reconstruction accuracy. The state-conditioned responses therefore reflect a mixture of genuine coordinated population fluctuations and a bias introduced by the analysis. Consequently, finding a partition in which the state-conditioned average responses separately reconstruct the conscious and suppressed faces does not establish that the population undergoes coordinated switching between these two representations.

### Coordinated population states alternate between conscious and suppressed stimulus representations

We therefore tested whether population activity exhibits coordinated transitions between representations of conscious and suppressed stimuli. We asked first whether a two-state model captured population activity better than the best-fitting single-state model, obtained by maximum likelihood estimation (see Methods). The two-state model assigned higher likelihood to the observed population activity than the best single-state model for all 19 sessions (**Fig. 3a**, *p* = 10^−4^, t-test). This indicates that the observed switching between two states reflects coordinated population dynamics rather than independent fluctuations of individual neurons.

**Figure 3.**
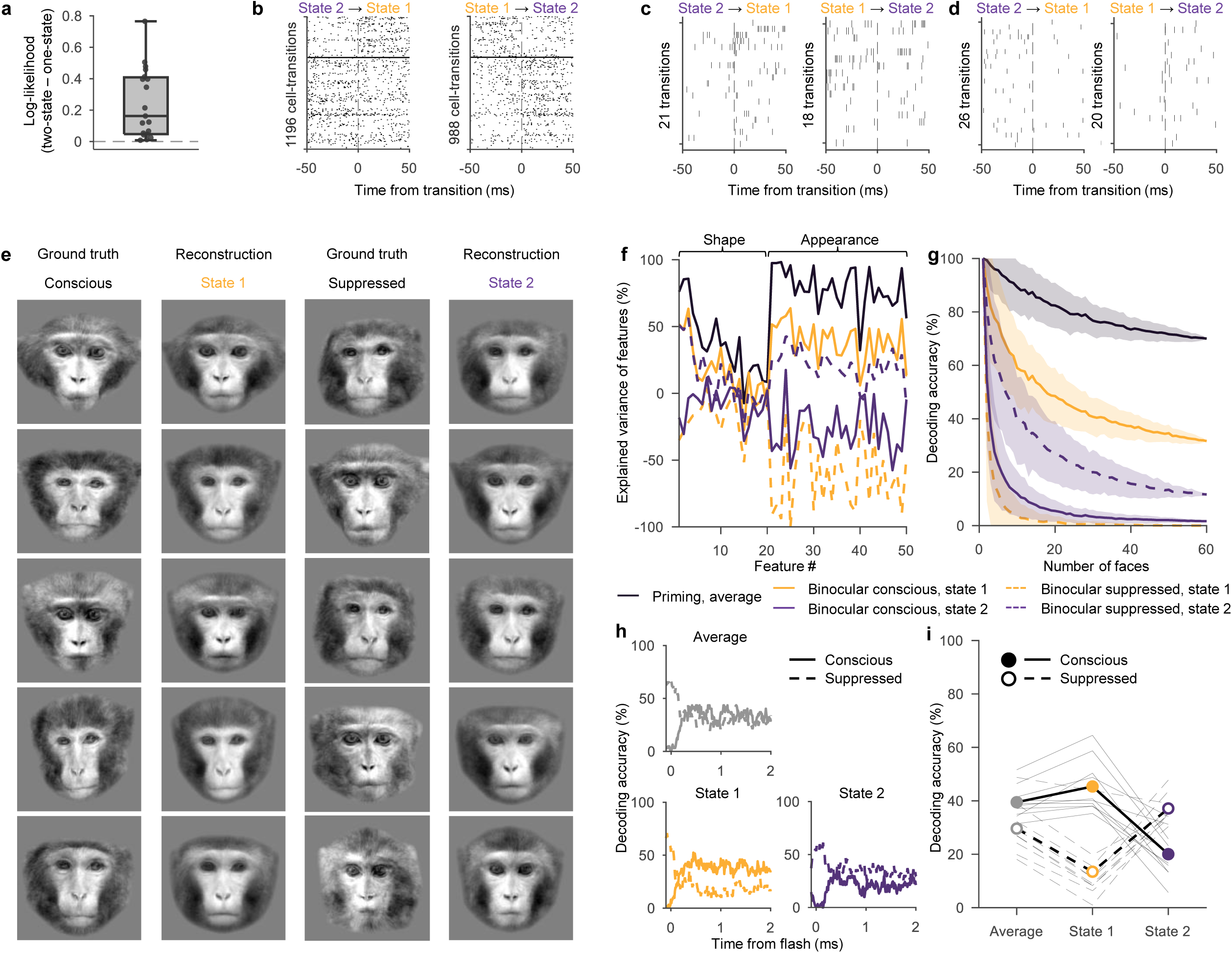
Coordinated switching of the population between conscious and suppressed states. (**a**) Difference of marginal log-likelihoods of two-state and one-state models. We computed per-trial, per-neuron gain in log-likelihood of the two-state model over the best-fitting single-state model, whose maximum-likelihood static firing rate is given by the average firing rate during binocular rivalry. Boxes, median and interquartile range across the 19 sessions; whiskers, range; points, individual sessions; horizontal dashed line indicates where neither model has an advantage. We compared the log-likelihood difference against zero using a two-sided t-test (*t*(18) = 4.84, *p* = 10^−4^, *n* = 19 sessions). (**b**) Same as Fig. 2e but for a held-out randomly selected half of neurons; the other half was used to infer states. (**c**)-(**d**) Same example cells as in Fig. 2f, g but all other neurons were used for inferring states. (**e**) Reconstructions as in Fig. 2h but using held-out half of neurons. (**f**) Explained variance as in Fig. 2i but using held-out half of neurons (t-test, *t*(29) = 17.5, *p* = 6 × 10^−17^, *n* = 30 stimulus pairs). (**g**) Decoding accuracy as in Fig. 2j but using held-out half of neurons (two-sided sign-flip permutation test, *p* = 10^−5^, *n* = 30 stimulus pairs). (**h**) Single-trial decoding for example session as in Fig. 2k but using held-out half of neurons. (**i**) Single-trial decoding summary as in Fig. 2l but using held-out half of neurons (two-sided paired t-test; state 1: *t*(9) = 7.6, *p* = 3 × 10^−5^, *n* = 10 sessions; state 2: *t*(9) = −6.5, *p* = 1 × 10^−4^, *n* = 10 sessions).

We next tested whether the representational content alternated in a coordinated manner across the population. We split neurons into two random halves, using one half of the population to infer the state sequence, and the other half to predict face features. If the inferred state sequence captured a genuine coordinated alternation, the neurons held out for prediction should change their represented stimulus when alternating between inferred states.

Rasters of populations of held-out neurons and individual held-out neurons showed the same changes in activity at inferred state transitions (**Fig. 3b-d**). Most importantly, the held-out population reconstructed the conscious face during epochs classified as state 1 by the other half of the neurons, and the suppressed face during epochs classified as state 2 (**Fig. 3e–i)**. Reconstructions using the held-out half of neurons were modestly less accurate than those obtained from the full population (as expected given the circularity bias described above) but continued to clearly separate conscious and suppressed face representations during state 1 and state 2 (*p* = 6 × 10^−17^, t-test, see Methods). The cross-validated results held true on an individual monkey level (**Extended Data Fig. 3**) and an individual session level (**Extended Data Fig. 4**). Together, these results demonstrate that the population genuinely alternates between conscious and suppressed representations.

We confirmed that the lower reconstruction accuracy for the held-out half was not due to using fewer neurons for state inference and face reconstructions. To do so, we inferred states from all but one neuron, used these states to compute the state-conditioned responses of the held-out neuron, and then pooled these responses across all held-out neurons to generate reconstructions (leave-one-neuron-out). This analysis removes circular conditioning without sacrificing population size. The leave-one-neuron-out validation increased face explained variance by only around 2 percentage points relative to cross-validation using half of neurons, showing that state inference and reconstruction accuracy were not limited by neuron count (**Extended Data Fig. 5a–c**). On the other hand, restricting the split-half analysis to high-confidence time points increased explained variance by 10 percentage points (**Extended Data Fig. 5d–f**), showing that these moments contained high-fidelity representations of conscious and suppressed stimuli.

### Control analyses confirm coordinated within-trial state alternation

Having established that the population exhibits coordinated dynamics, we next controlled for two alternative hypotheses: the first alternative is that the states reflect differences between trials rather than within-trial switching; the second alternative is that interleaved representations of conscious and suppressed stimuli are imposed by the inference procedure itself.

We first asked whether the observed alternation between conscious and suppressed stimuli reflected genuine *within-trial* switching or rather differences between trials. Even though the analysis was restricted to the 700 ms period in which human observers reliably perceive the flashed face, the two states could in principle arise from some trials being dominated by the conscious face and others by the suppressed face, rather than from switching within each trial. To rule this out, we subsampled time points within each trial so that every trial contributed equal numbers of samples to state 1 and state 2. This ensured that the two state-resolved reconstructions were based on matched numbers of samples from the same trials, eliminating differences between trials as a source of the state separation. Despite the fact that this procedure discarded many time points, state-resolved reconstructions remained robust (**Extended Data Figure 6**). Thus, the separation between conscious and suppressed representations cannot be explained by different stimuli being represented on different trials; instead, both representations are expressed within single trials in alternating micro-epochs.

Second, we randomly shuffled the response templates used to define the two states and repeated the reconstruction analysis. This tested whether our state-inference procedure could manufacture accurate reconstructions of any stimulus simply by sorting time points to whatever templates it was given. If it could, this procedure should yield reconstructions that match those shuffled templates even though they were not presented while recording the activity used for reconstruction. To the contrary, reconstructions obtained with randomly re-assigned templates did not resemble the templates, and their explained variance was largely below zero (**Extended Data Fig. 7a–c**). We also compared reconstructions obtained with shuffled templates to the faces that were actually presented on each trial (**Extended Data Fig. 7d–f**). These reconstructions were much worse than those obtained with the correct templates, indicating that accurate reconstruction depends on the template used for state inference matching the true face pair, and does not arise from arbitrary partitions of neural activity.

### Single cells have higher gain of tuning during conscious compared to suppressed epochs

The reconstruction results, together with the control analyses above, establish that the population alternates between representing the conscious and suppressed face. This raises the question of how the two representations differ at the single-cell level, and whether the conscious representation carries any signature that distinguishes it from the suppressed one. We therefore asked how individual face cells were tuned to each face, and how that tuning changed between the two states. As in **Fig. 3**, we inferred states based on one half of neurons, and measured single-cell tuning in the held-out half of neurons.

Face cells exhibit axis tuning: they respond as a linear function of a face’s position along a preferred axis in shape-appearance space^14^. If this tuning is preserved during rivalry, a cell’s responses should ramp along its preferred axis as the represented face varies. To test this, we ordered the 1500 binocular stimuli by the response each cell was predicted to give to one of the two faces, using weights fit to priming responses, and plotted the cell’s actual binocular responses in that order; a cell representing that face should produce responses that ramp along the ordering of stimuli. We performed analyses separately for orderings by the conscious and the suppressed face, and separately for time points assigned to state 1 and state 2.

In individual cells, the ramp appeared for the conscious face during state 1 and for the suppressed face during state 2 (example cell in **Fig. 4a**). During state 1, a cell’s responses increased steadily as the conscious face moved along its preferred axis, while ordering the same responses by the suppressed face produced no gradient. During state 2 this reversed: the same cell now ramped with the suppressed face and showed no gradient for the conscious face.

**Figure 4.**
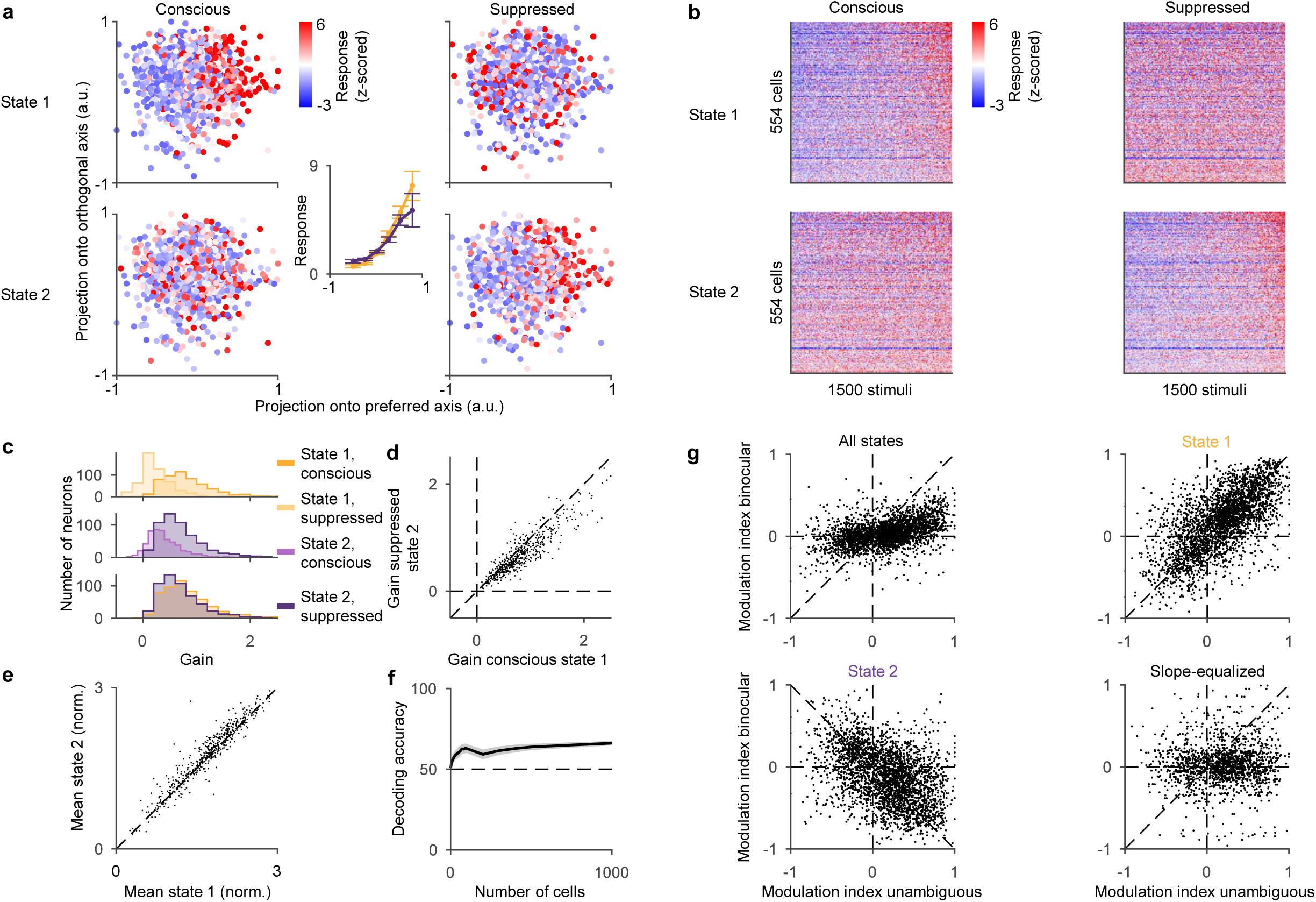
Single cells switch between conscious and suppressed representations. (**a**) Ramp-shaped tuning to features of conscious and suppressed faces during state 1 and state 2 for an example cell. Each dot represents one of the 1500 images, projected onto the cell’s preferred axis (x-axis) and principal orthogonal axis (y-axis). The colour of each dot represents the cell’s response to the image. Dot positions are determined by the features of the conscious face (left) or the suppressed face (right). Responses were computed during state 1 (top) or state 2 (bottom). The center inset shows marginal distributions for the conscious face in state 1 (orange) and suppressed face in state 2 (purple) as a function of the projection onto the preferred axis. Since red and cyan faces were presented through the anaglyph goggles, we removed the first principal component of features to remove this colour component for (a)-(f). The results without removing this component were qualitatively the same. (**b**) Population tuning across all cells to conscious face features (left) and suppressed face features (right) during state 1 (top) and during state 2 (bottom). For each cell (rows of each subpanel), colour indicates responses to each of the 1500 stimuli (columns), sorted by the projection onto the preferred axis of each cell (inferred by linear regression during priming). Cells are sorted by the slope of their response gradient, which we refer to as tuning gain. (**c**) Histogram of tuning gain of single cells for the features of the conscious face during state 1 (orange), features of the suppressed face during state 2 (purple), features of the suppressed face during state 1 (bright orange), and features of the conscious face during state 2 (bright purple). We compared gains for conscious and suppressed using a two-sided paired t-test (state 1: *t*(542) = 25.5, *p* = 8 × 10^−95^, *n* = 543 cells; state 2: *t*(542) = −18.6, *p* = 6 × 10^−60^, *n* = 543 cells). (**d**) Sharper tuning for conscious features during state 1. Each dot represents one cell. X-axis corresponds to gains for features of the conscious face during state 1. Y-axis corresponds to gains for features of the suppressed face during state 2. Dashed diagonal line indicates where the two gains are equal. We evaluated the difference between the gains with a paired t-test (*t*(542) = 14.4, *p* = 3 × 10^−40^, *n* = 543 cells) and computed the correlation between gains (Pearson’s *r* = 0.90, *p* = 5 × 10^−192^, *n* = 543 cells). (**e**) Slightly stronger mean responses during state 2. Same conventions as (d), but for mean response across stimuli instead of gain. We evaluated the difference with a two-sided paired t-test (*t*(542) = −8.0, *p* = 2 × 10^−15^, *n* = 543 cells). (**f**) Decoding accuracy for discriminating state 1 vs state 2 across all 1500 stimuli. A linear decoder used the firing rate of all neurons. Shaded regions indicate standard deviation across 100 decoding iterations with randomly selected subset of cells and random 50/50 split of stimulus pairs into train and test set. Dashed line indicates 50% chance level. (**g**) Unambiguous modulation vs perceptual modulation during average activity (top left; Pearson’s *r* = 0.45, *p* = 7 × 10^−147^, *n* = 2885 cell-stimulus pairs), state 1 (top right; Pearson’s *r* = 0.66, *p* = 2 × 10^−355^, *n* = 2885 cell-stimulus pairs), state 2 (bottom left; Pearson’s *r* = −0.44, *p* = 5 × 10^−139^, *n* = 2885 cell-stimulus pairs), and activity averaged across gain-normalized state 1 and state 2 responses (bottom right; Pearson’s *r* = −0.02, *p* = 0.3, *n* = 2885 cell-stimulus pairs; see Methods). As in Fig. 3, states were inferred using one half of neurons, and analysis was performed using the held-out half of neurons.

This pattern held across the population (**Fig. 4b**). A clear gradient was present for the conscious face during state 1 and for the suppressed face during state 2, with little gradient for the off-target face in each state. Rivalry therefore preserved each cell’s underlying feature tuning but expressed it selectively for the face represented by the population at a given moment. Confirming this, the cell’s preferred tuning axis computed for the rivalry responses to the conscious face during state 1 matched the tuning axis computed from priming responses, barely lower than noise ceiling (**Extended Data Figure 8;** *p* = 0.01, paired t-test). Similarly, the tuning axis from responses to the suppressed face during state 2 was indistinguishable from the priming tuning axis relative to noise ceiling (*p* = 0.35). On the other hand, responses to the suppressed face during state 1, and the conscious face during state 2 did not yield tuning axes consistent with the priming tuning axis (state 1 suppressed: *p* = 10^−87^, state 2 conscious: *p* = 6 × 10^−69^).

Comparing gain of tuning, measured by the slope of the response against the preferred feature axis, confirmed that during state 1, tuning to the conscious face was significantly steeper than tuning to the suppressed face (**Fig. 4c**, top; *p* = 8 × 10^−95^, paired t-test); during state 2, the reverse held, with tuning to the suppressed face significantly steeper than to the conscious face (**Fig. 4c**, middle; *p* = 6 × 10^−60^, paired t-test).

Despite this apparent symmetry, the two states were not mirror images of each other. The gain of tuning to the conscious face during state 1 was higher than that to the suppressed face during state 2 (**Fig. 4c**, bottom; *p* = 3 × 10^−40^, paired t-test; mean ratio between conscious and suppressed tuning gain = 1.21). Across cells, tuning gain for the conscious face during state 1 was strongly correlated with gain for the suppressed face during state 2 (**Fig. 4d**, Pearson’s *r* = 0.90), indicating that the same neurons encoded both faces. We also compared the mean responses across all stimuli during state 1 and state 2, and found that, unlike tuning gain, the mean firing rate was weakly but significantly higher during state 2 for most neurons (**Fig. 4e**, *p* = 2 × 10^−15^, paired t-test). The two states therefore differ in two respects: a tuning-gain difference (steeper in the conscious state) and a small mean-rate offset (higher in state 2). A linear decoder distinguished state-1 from state-2 responses with 66% accuracy (**Fig. 4f**, black, see Methods).

### Temporal multiplexing explains weak average modulation during binocular rivalry

We next revisited the original puzzle, why perceptual modulation during binocular rivalry appears weak when averaging across time (**Fig. 4g**, top left). One possibility is that during state 1, cells are strongly modulated by the conscious stimulus, and during state 2 cells are strongly modulated by the suppressed stimulus, and averaging across both states yields apparent weak modulation. To test this, we computed modulation indices separately for time points assigned to state 1 and state 2, inferred from a held-out half of neurons as in **Fig. 3**.

Strikingly, we found that modulation by conscious percept in the binocular condition during state 1 was approximately as strong as modulation for average unambiguous responses (**Fig. 4g**, top right, Pearson’s *r* = 0.66). Modulation indices clustered around the diagonal (slope = 0.93), indicating that during state 1, cells responded as if the conscious stimulus was presented unambiguously. Conversely, during state 2, the pattern reversed, and modulation by conscious percept in the binocular condition was strongly anticorrelated with average unambiguous responses (**Fig. 4g**, bottom left, Pearson’s *r* = −0.44). Modulation indices clustered around the antidiagonal (slope = −0.76), indicating that during state 2, cells responded as if the suppressed stimulus was presented unambiguously.

These results suggest that the weak average modulation was observed because averaging collapses responses across the two opposing states: state 1, which represents the conscious stimulus, and state 2, which represents the suppressed stimulus. However, this explanation raises a further question: *why does any positive modulation remain after averaging across the two states?* In the previous section we found that the gain of tuning to conscious stimuli during state 1 was slightly higher than the gain to suppressed stimuli during state 2. We next asked whether this asymmetry could account for the residual positive correlation between binocular and unambiguous modulation indices.

To test this, we re-computed modulation indices for responses where we equalized the tuning gain for state 1 and state 2, and averaged the gain-equalized responses, weighted by the occupancy of state 1 and state 2 for each stimulus pair (**Fig. 4g**, bottom right; see Methods). Interestingly, after this correction the residual positive correlation disappeared (Pearson’s *r* = −0.02, *p* = 0.31; slope = 0.07). Thus, weak average modulation during binocular rivalry can be explained by averaging across two temporally interleaved states with opposite stimulus selectivity, combined with slightly stronger tuning in state 1.

### Dynamics of state switching

We next asked how fast the population switches between the two states, and how long it dwells in each. Our earlier analysis labelled each time bin independently, ignoring any dependence between successive bins. To ask whether the states persist over time, and if so, for how long, we turned to a Hidden Markov Model (HMM), fit separately for each stimulus pair. As in the earlier LLR analysis, the two states emitted spikes from fixed templates (the pair’s priming-derived firing rates for the conscious and suppressed face), with the additional assumption that transitions between states were governed by a Markov process; only the 2×2 transition probability matrix was free, and it was estimated from the data (**Fig. 5a**). Fitting the model to that pair’s binocular trials inferred the transition probabilities and the state sequence that maximized the probability of the observed spike counts, thereby partitioning each trial into two states over time (**Fig. 5b**). To determine the characteristic time scale of state alternations, we measured the median dwell times of the inferred states. Because dwell time cannot be resolved below the width of a single bin, it is expected to increase with bin size once the bin width becomes sufficiently large, and this dependence must be accounted for before the estimate can be interpreted. We therefore fit the HMM across a range of bin sizes and, at each bin size, asked how well the inferred state sequence explained the spikes of held-out neurons, relative to a one-state model (normalized conditional log-likelihood ratio; Methods). This measure peaked at small bin sizes, and over this range median dwell time plateaued around ∼60 ms (**Fig. 5c**), indicating that the estimate reflects the data rather than the temporal resolution of the analysis. Specifically, for the peak-likelihood bin size of 5 ms, median dwell times for state 1 and state 2 were very similar, 64 ms and 59 ms, respectively (**Fig. 5d**). This is consistent with the 67 ms time constant of likelihood autocorrelation reported earlier (**Fig. 2d**).

**Figure 5.**
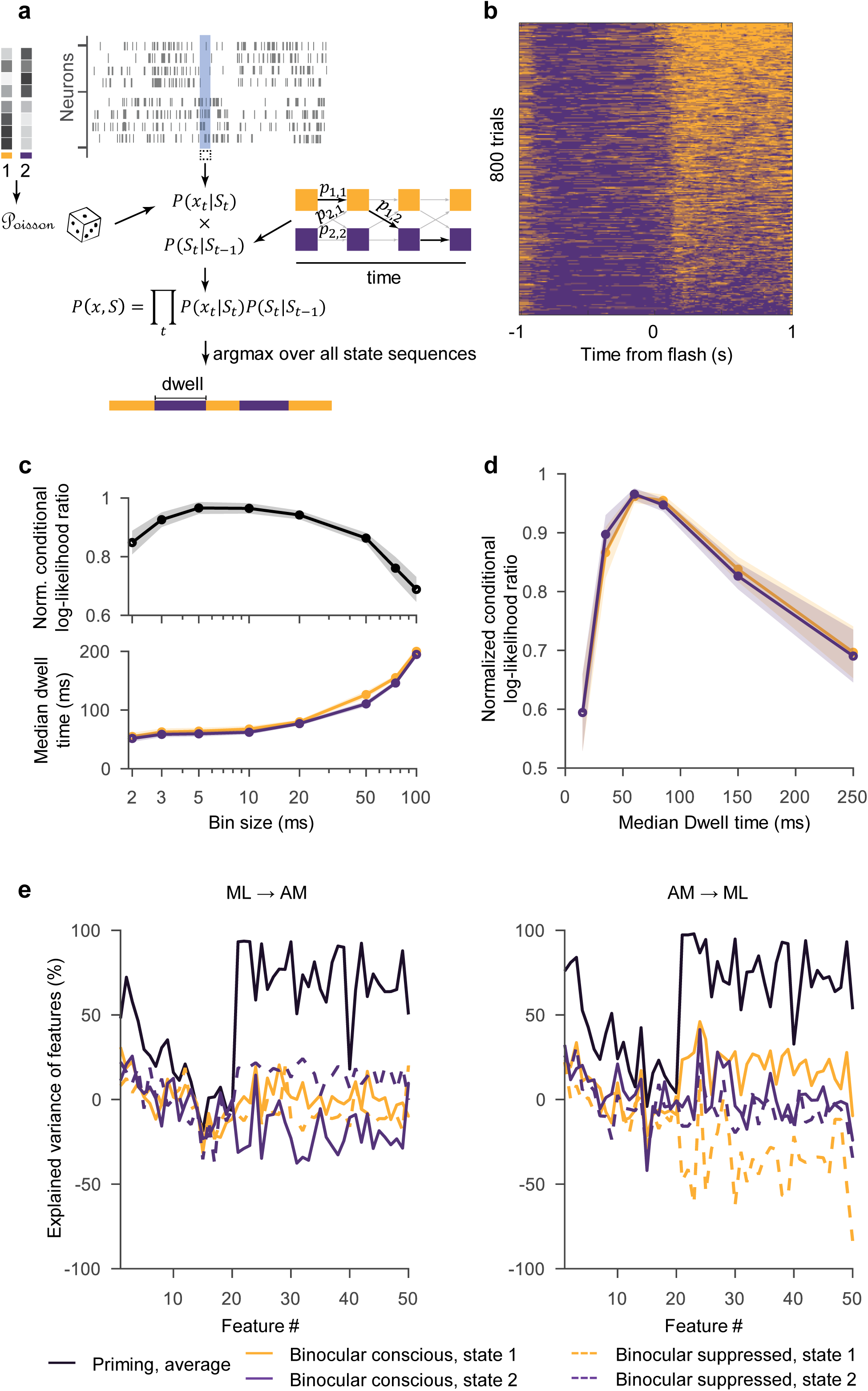
Dynamics of state alternations. (**a**) Schematic of Hidden Markov Model (HMM). The HMM finds the most likely partition of the time course into two states given inferred state transition probabilities and the observed spike counts. (**b**) State partitions of all trials (rows), sorted by proportion of state 1, for an example session. (**c**) Fit accuracy (top) and median state dwell times (bottom) for state 1 (orange) and state 2 (purple) as a function of bin size parameter. Fit accuracy is quantified using a conditional log-likelihood ratio of a two-state HMM and a one-state HMM, normalized by its peak value across bin sizes per session. The emission vector in the one-state HMM was fixed at the mean of the two templates. Shaded regions indicate s.e.m. across sessions. (**d**) Fit accuracy (y-axis) as a function of median dwell time (x-axis, binned). Shaded regions indicate s.e.m. across sessions. (**e**) Explained variance of face-feature reconstruction when the two HMM states are inferred in one area and the features are reconstructed from the other. Left, states inferred from ML, reconstructed from AM; right, states inferred from AM, reconstructed from ML (paired t-test; ML→AM: *t*(29) = 4.1, *p* = 3 × 10^−4^, *n* = 30 stimulus pairs; AM→ML: *t*(29) = 5.4, *p* = 9 × 10^−6^, *n* = 30 stimulus pairs). Same conventions as Fig. 2i. Only 7 of the 9 sessions were included because cross-region analyses required simultaneous recordings from both ML and AM.

### On and Off states

Single-trial population activity showed strong fluctuations in overall firing, resembling previously described On and Off states^16^ (**Fig. 1g**). Details regarding these On-Off dynamics within face patches will be the focus of a separate forthcoming manuscript (C.M., J.K.H., F.L.L., D.Y.T., T.A.E., in preparation). Here, we address the specific concern that our two states might track this gain rhythm (high versus low excitability) rather than which stimulus is represented (conscious versus suppressed). To test this directly, we used a modified version of our log-likelihood ratio decoder that used two template vectors for each stimulus, one for the high-gain On periods and another for the low-gain Off periods. This four-state LLR decoder, like the original two-state LLR decoder, labelled each time point with the template vector that maximized the likelihood of the observed spikes. The On-Off axis absorbed the large excitability fluctuations, yet the conscious-suppressed axis remained independent of it: (i) conscious and suppressed states were equally represented within both gain regimes (49%/51% during On, 50%/50% during Off), (ii) the conscious/suppressed partition agreed with a partition by total firing rate (separating bins into higher/lower than median firing rate) no better than chance (Cohen’s κ ≈ 0.0), and (iii) the conscious/suppressed states recovered by the four-state model agreed closely with those from the original two-state model (**Fig. 2a**; 80% agreement), leaving the state-resolved reconstructions and single-cell tuning asymmetry largely unchanged (**Extended Data Figure 9**). The On-Off fluctuations therefore did not substantially affect the inference of conscious/suppressed states.

### Alternations of conscious and suppressed states are weakly coordinated across face patches

At the perceptual level, rivalry is unified: only one image is consciously seen at a time, and the corresponding representation is thought to be coherent across the brain. It is not clear, however, that this unity extends to the fast micro-epochs uncovered here. The alternation between conscious and suppressed states could be a coordinated, network-wide event, with regions switching together, or each region could alternate on its own clock, arriving at the same percept through independent selection. To distinguish these possibilities, we inferred the state sequence separately from the simultaneously recorded ML and AM populations, each with its own neurons and its own templates (**Fig. 2a**), and then predicted face features using neurons in the other patch (**Fig. 5e**). Explained variance of features was relatively low compared to **Fig. 3**, and for AM→ML, explained variance for the conscious face in state 2 was higher than that for the suppressed face; combining across conditions, we did find a significant separation overall between conscious representations in state 1 and suppressed representations in state 2 (ML→AM: *p* = 3 × 10^−4^; AM→ML: *p* = 9 × 10^−6^, t-test, see Methods). However, our results do not provide strong evidence for coordination across patches in generating conscious and suppressed micro-epochs.

### Comparison of two-state model to alternative models

Several alternative mechanisms could in principle reproduce one or another signature of our analyses so far without reflecting a genuine, population-level temporal multiplexing of conscious and suppressed representations. We therefore considered a broad set of alternative models and subjected each, together with the recorded data, to a battery of complementary control analyses (**Extended Data Fig. 10a**, model × test matrix). Only a two-state model, in which the same neurons coherently alternate, within single trials, between faithfully representing the conscious and the suppressed face, reproduced every signature observed in the real data; each alternative failed at least three of the tests.

#### Alternative models

We simulated four generative models of single-trial neural activity, each built from the same priming-derived, per-neuron response templates for the two faces of a trial (*r_A_* conscious, *r_B_* suppressed) and matched to the recorded trial structure of the real data (see Methods): (i) a two-state model, in which the entire population coherently alternates between the two faces at the ∼60 ms timescale estimated from the data; (ii) a conscious-only model in which only the conscious face is represented throughout the trial; (iii) an averaging model, in which every neuron fires at a fixed rate equal to the average of its two priming templates, with no alternation; and (iv) an uncoordinated model, in which every neuron alternates between the conscious and suppressed face with the same ∼60 ms dwell time as the two-state model but on its own, independent of the rest of the population. In every model, spike counts were generated as Poisson draws from the model’s per-neuron, per-time-bin firing rate.

#### A battery of tests

We evaluated every model, and the real data, on four tests (matrix rows in **Extended Data Fig. 10a**): (i) reconstruction of the conscious and suppressed faces from state-resolved activity using circular analysis (as in **Fig. 2h,i**); (ii) coordinated reconstruction, the same reconstruction after the state is inferred from one half of the population and the faces are reconstructed from the held-out half (as in **Fig. 3**); (iii) reversal, the same tuning reversal measured in held-out neurons (as in **Fig. 4g**); (iv) model comparison of whether a two-state description of the activity fits held-out data better than a one-state alternative. In all cases, we used an HMM for state partitioning.

The conscious-only model cannot reconstruct the suppressed face for obvious reasons, and therefore is clearly incompatible with our experimental findings; we therefore focus on the three models that represent both faces. The two-state, averaging, and uncoordinated models could all reconstruct the conscious face in inferred state 1 and the suppressed face in inferred state 2 above chance using circular analysis (**Extended Data Fig. 10b**). This may at first seem surprising: how could a model with no genuine alternation, e.g., an averaging model that never represents either face on its own, be partitioned into two sets of time points from which noise alone reconstructs each pure face? Yet this is actually expected on deeper examination. In the averaging model every neuron’s response sits at the mean of its two responses, 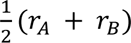, and only Poisson noise makes the activity vary; the HMM assigns the moments that happen to deviate toward *r_A_* to state 1 and those toward *r_B_* to state 2. Since the same data samples used for state inference are also used for computing state-conditioned responses, reconstructing from these selected moments then recovers *r_A_* and *r_B_* due to circular conditioning. Any model that contains both faces’ responses—averaged together (averaging model), or switched independently (uncoordinated)—passes this test for the same reason. However, note that the averaging model, though seemingly simple, is actually quite unnatural: representing two competing stimuli as the average of their individual responses is a linear-superposition property that a strongly nonlinear system such as the ventral visual stream would not generally exhibit. Importantly, inferred states switched much more frequently for the averaging model data than for the real data, allowing for exacerbated circular conditioning; when matching dwell times (see Methods), real data yielded significantly better reconstructions than the averaging model (Δ = 8.2% explained variance at ∼80 ms dwell, *p* = 0.007, bootstrap over n = 60 stimulus pairs, **Extended Data Fig. 10c**). Still, above-chance state-resolved reconstruction is expected even of models with no genuine alternation, and on its own is not diagnostic of a genuine, coordinated alternation.

The decisive question is whether these fluctuations are coordinated across neurons rather than private to each cell. When the state was inferred from one half of the population and the conscious versus suppressed face features reconstructed from the held-out half (**Extended Data Fig. 10d**), reconstruction held in the real data and the two-state model but collapsed for the averaging and uncoordinated models, whose apparent alternation arises from independent single-neuron fluctuations that are not shared across neurons.

The lower cross-validated explained variance in the real data compared with the idealized two-state simulation (**Extended Data Fig. 10b,d**) may partly reflect the fact that transitions in biological neural populations are unlikely to be instantaneous. While the simulated activity jumps directly between two discrete states, real population activity is expected to take a finite time to move between locations in neural state space, producing intermediate activity patterns. Forcing these transition periods into one of two states may reduce state-specific reconstruction accuracy and hence explained variance.

Single-cell tuning showed the same pattern as above: When inferring the state from one half of the population and measuring tuning in the held-out half, only the real data and the two-state model retained a reversal, whereas the averaging and uncoordinated models collapsed to zero (**Extended Data Fig. 10e**).

As a complement to the preceding tests, we asked whether the population activity is statistically better described by two states than by one. For the real data and for each model, we fit two-state and single-state HMMs and compared their conditional likelihoods on held-out data (**Extended Data Fig. 10f**). For the single-state model the one rate was a weighted average of the two faces’ priming rates; we scanned the averaging weight from 0 to 1 and kept the maximum-likelihood value. Two states fit the real data substantially better than one, establishing that the activity is genuinely organized around two population states; the coordinated two-state model showed the same advantage. For the averaging and uncoordinated models, by contrast, a single state fit better than two: adding a second state captured only noise and reduced the held-out likelihood.

### Evidence for similar interleaved representations in prefrontal cortex

The results so far show that inferotemporal cortex interleaves representations of the conscious and suppressed face. We next asked whether prefrontal cortex carries a pure representation of the conscious face, or an interleaved representation similar to inferotemporal cortex. We repeated the flash-suppression experiments in prefrontal cortex, recording from the lateral orbitofrontal sulcus and the infraprincipal dimple (**Fig. 1e, right**).

Reconstructions were of lower quality than in inferotemporal cortex (**Fig. 6a**): the best possible explained variance, reconstructing faces from priming responses, was below 80%, compared to over 90% for inferotemporal cortex, and particularly low (less than 30%) for shape features (**Fig. 6b**). This may reflect a lower proportion of face cells or a different feature code in prefrontal cortex. Despite this, the same pattern of state-dependent representation of conscious and suppressed stimuli held in prefrontal cortex: reconstructions during state 1 more closely resembled the conscious face, and those during state 2 the suppressed face, confirmed by both explained variance and decoding using the same data samples as for state inference (**Fig. 6b-c**, *p* = 10^−14^, t-test, see Methods). Importantly, as in inferotemporal cortex, the effect held under a cross-neuron validation: when the states were inferred from one half of the prefrontal population and the faces reconstructed from the held-out half, the state-dependent separation persisted (**Fig. 6d-e**, *p* = 3 × 10^−4^, t-test, see Methods), indicating a genuine population-level alternation.

**Figure 6.**
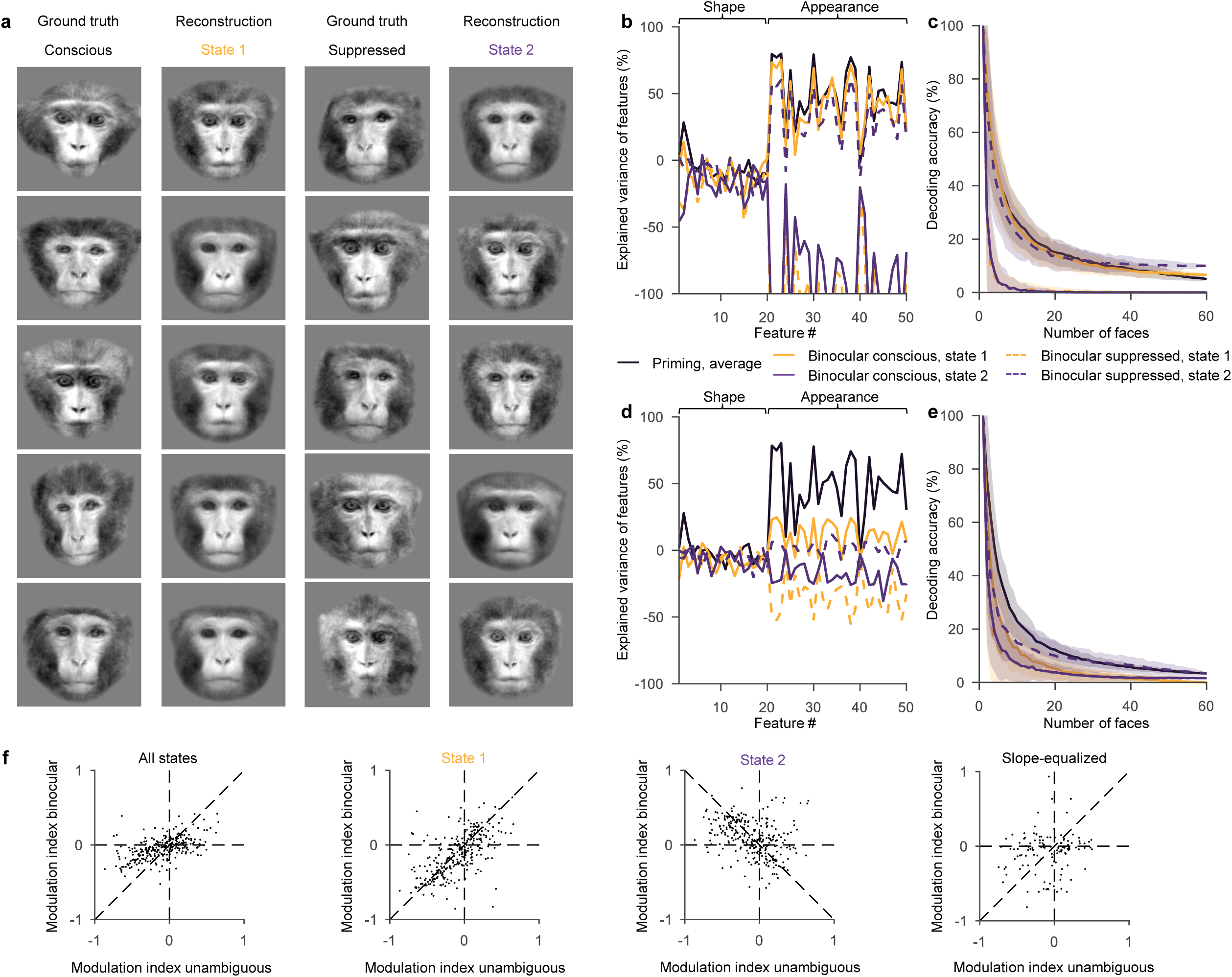
Temporal multiplexing in prefrontal cortex. (**a**) Reconstructions of faces during state 1 and state 2 for prefrontal cortex. Same conventions as Fig. 2h but LLR templates were computed and features predicted from neural activity in prefrontal patches PO and PVL. (**b**) Explained variance of features. Same as Fig. 2i but for PFC (t-test, *t*(29) = 14.2, *p* = 10^−14^, *n* = 30 stimulus pairs). (**c**) Decoding accuracy. Same as Fig. 2j but for PFC (two-sided sign-flip permutation test, *p* = 10^−5^, *n* = 30 stimulus pairs). (**d**) Explained variance of features using held-out half of neurons as in Fig. 3f, but for PFC (paired t-test, *t*(29) = 4.1, *p* = 3 × 10^−4^, *n* = 30 stimulus pairs). (**e**) Decoding accuracy, using held-out half of neurons as in Fig. 3g, but for PFC (two-sided sign-flip permutation test, *p* = 0.03, *n* = 30 stimulus pairs). (**f**) Unambiguous modulation vs perceptual modulation. Same as Fig. 4g but for PFC. We compared average perceptual modulation to unambiguous modulation using a two-sided paired t-test (*t*(282) = −3.5, *p* = 5 × 10^−4^, *n* = 283 cell-stimulus pairs). Correlation between unambiguous and perceptual modulation indices was quantified using Pearson’s correlation (*r* = 0.56, *p* = 3 × 10^−24^; *n* = 283 cell-stimulus pairs for average, *r* = 0.63, *p* = 1 × 10^−32^; *n* = 283 cell-stimulus pairs for state 1, *r* = −0.32, *p* = 5 × 10^−8^; *n* = 283 cell-stimulus pairs for state 2, *r* = 0.24, *p* = 0.005; *n* = 283 cell-stimulus pairs for gain-equalized).

Modulation by conscious percept was again much weaker compared to unambiguous alternation for average activity (**Fig. 6f**, *p* = 4 × 10^−10^, paired t-test; slope = 0.32), suggesting that, just like inferotemporal cortex, prefrontal cortex also does not have a pure representation of conscious perception for average responses. During state 1, however, modulation resembled the conscious stimulus (slope = 0.91) and during state 2 modulation resembled the suppressed stimulus (slope = −0.52). Temporal multiplexing of conscious and suppressed representations may therefore be a broad cortical phenomenon rather than a feature unique to inferotemporal cortex.

## Discussion

The findings presented above reveal a temporally multiplexed code for conscious and suppressed visual representations. State-resolved reconstructions show that inferotemporal cortex alternates between complementary representational epochs: one in which the population represents the consciously perceived face, and another in which it represents the suppressed face. These representational switches in neural activity happen over an order of magnitude faster (15 Hz) than perceptual switches (∼0.5 Hz)^6,17^. Single-cell analyses show that this alternation is not produced by separate populations of conscious- and suppressed-selective neurons, but by the same neurons changing which stimulus they represent across states. Importantly, during the conscious state, single-cell tuning to the conscious stimulus has stronger gain than tuning to the suppressed stimulus in the suppressed state, and a linear decoder can discriminate state 1 and state 2 across stimulus pairs. These states are weakly coordinated across simultaneously recorded face patches. Finally, prefrontal cortex shows a qualitatively similar interleaved structure. Together, these findings show that time provides a critical dimension for separating conscious and suppressed representations within cortical population activity.

Our findings expose a limitation of searching for the neural correlates of consciousness within the average activity of specific brain regions, whether toward the front or back of the brain. Rather than finding a pure representation of conscious perception in high-level areas, we found that average rivalry responses remained less modulated than responses to unambiguous stimulus alternation, even in prefrontal cortex; this is consistent with previous studies^7^. The apparent weak modulation can now be understood as the consequence of averaging across temporally interleaved conscious and suppressed representations. The discovery of this temporal structure highlights the importance of large-scale simultaneous recordings for resolving coordinated population dynamics within single trials rather than relying on responses averaged across trials.

Previous studies noted that responses during rivalry did not fully collapse to the conscious stimulus response, leading the authors to suspect that suppressed stimuli might still leave residual information in neural activity^7^. This possibility was later supported by the finding that suppressed stimuli could be decoded from inferotemporal responses^2^. The present work extends these findings by showing that neural activity not only allows discriminating between two different suppressed stimuli but allows reconstruction of the suppressed stimulus with high fidelity. Thus, the suppressed stimulus is not represented as a weak residual trace but occupies the same high-dimensional face space as the conscious stimulus during complementary temporal epochs, and both epochs rely on a shared mapping between face features and single-cell responses. The presence of a suppressed stimulus representation even at high levels of the cortical hierarchy may help explain how unconscious stimuli are able to influence behaviour^18^, attention^19^, and affective responses^20^.

The temporal multiplexing reported here may reflect a broader coding principle by which neural populations organize information over time. In attention, behavioural sensitivity and frontoparietal activity fluctuate rhythmically, suggesting that visual information is sampled in alternating temporal windows rather than through a continuously stable spotlight^21–23^. In working memory, stimulus information can be expressed in brief beta- and gamma-band events, and recent work suggests that frontal theta phase can modulate which remembered location is most accessible for readout, indicating that temporal structure can regulate when stored representations are expressed or accessed^24–28^. In value-based decision-making, neural populations in orbitofrontal cortex alternate on a single-trial basis between states representing the values of different choice options^29^. In spatial navigation, abrupt changes in context can produce theta-paced flickering between distinct hippocampal place-cell maps^30^. Together with our findings, these studies suggest that temporal structure may be a general organizing principle for encoding multiple representations within overlapping neural populations.

The presence of rich representations of the suppressed stimulus raises a question for future work: why do representations in some micro-epochs reach awareness, whereas others do not? The fact that conscious and suppressed states can be discriminated by a linear decoder suggests that neural activity contains information not only about stimulus content, but also about whether the content represented in a given temporal epoch is destined for awareness or remains unconscious. One possibility is that this signature determines whether a sensory representation is dynamically routed to downstream systems involved in report, introspection, and behavioural control. Such routing could be controlled by recurrent interactions within visual cortex or by feedback from higher-order regions such as parietal cortex, the pulvinar, or other networks capable of coordinating perceptual states. Speculatively, suppressed epochs may allow the brain to explore alternative interpretations of the visual input, without being broadcast or influencing behaviour, whereas conscious epochs may express a working hypothesis that, through dynamic routing, becomes accessible to circuits that use it for introspection or actions such as report. Identifying the mechanisms that orchestrate these transitions, and determining why only some temporal segments become available to awareness, will be essential for understanding how temporal population dynamics give rise to conscious perception.

## Extended Data Figures

**Extended Data Figure 1.**
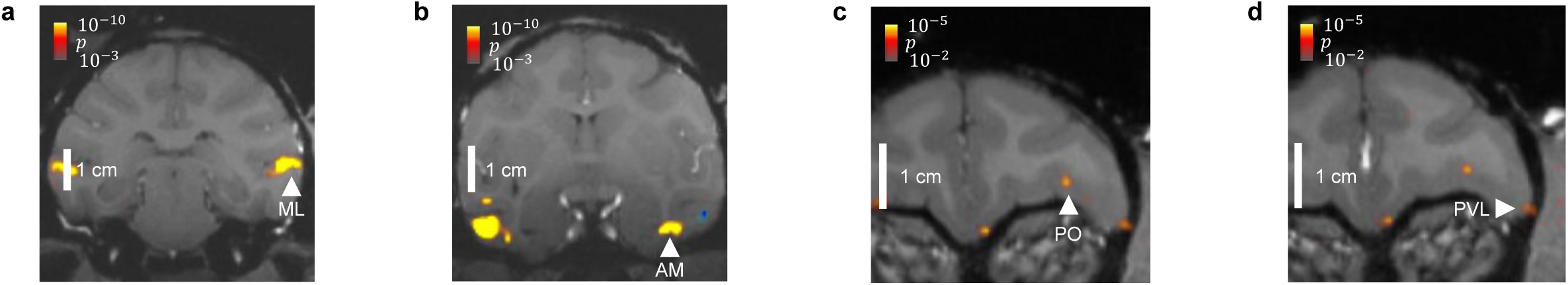
Recording targets. MRI anatomical scans showing targeted face patches in inferotemporal cortex (**a**) ML and (**b**) AM, as well as in prefrontal cortex (**c**) PO and (**d**) PVL. Colour overlay shows fMRI contrasts for visual activation to faces versus objects, which was used to guide targeting.

**Extended Data Figure 2.**
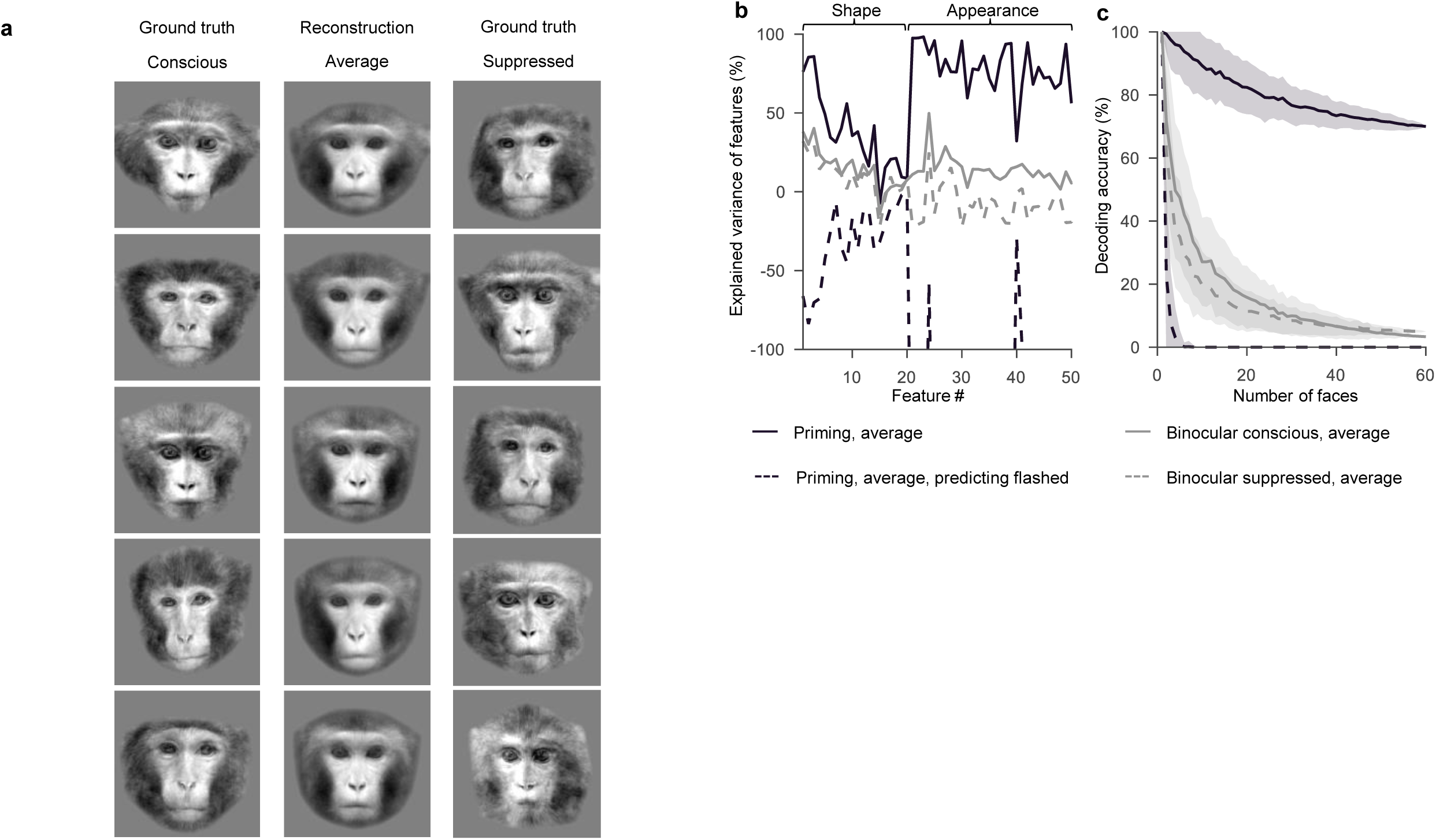
Weak reconstructions from average responses. (**a**) Reconstructions as in Fig. 2h but using average activity rather than state-resolved activity. (**b**) Explained variance of predicting features of the conscious (solid) or suppressed (dashed) face as in Fig. 2i but using average activity (grey). The black dashed line shows the similarity between the face reconstructed from priming activity and the other face of the stimulus pair, i.e., the face that will subsequently be presented as the flashed stimulus, but was not yet present during the analyzed priming period. As the face has not appeared yet during the analyzed period, this is a measure for chance performance. (**c**) Euclidean-distance decoding based on predicted features as in Fig. 2j but using average activity.

**Extended Data Figure 3.**
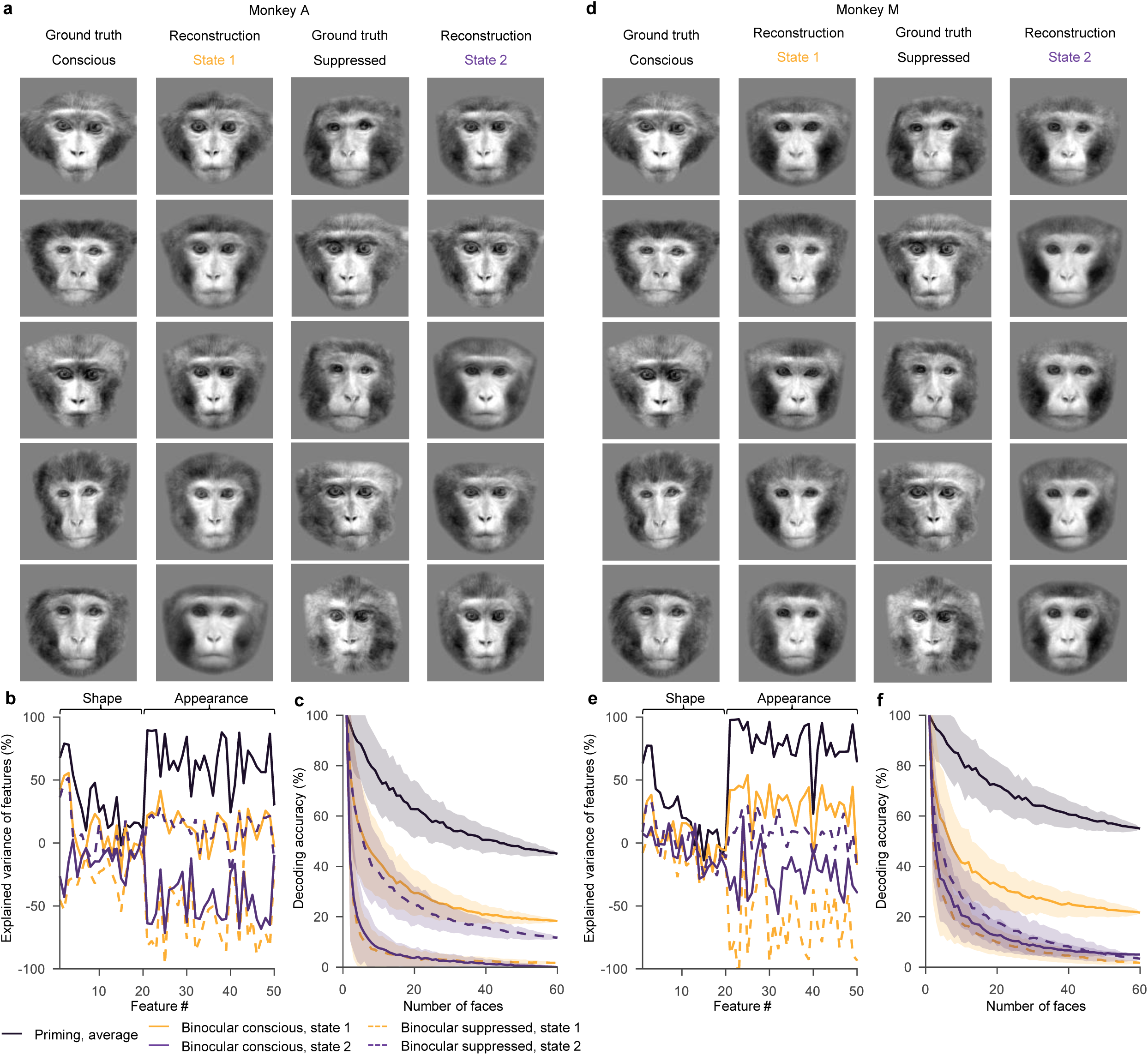
Interleaved representations in individual monkeys. (**a**)-(**c**) Same as Fig. 3e-g but pooling only across sessions from monkey A. (**d**)-(**f**) Same as Fig. 3e-g but pooling only across sessions from monkey M. As in Fig. 3, states were inferred from one half of neurons and features were predicted using the held-out half of neurons.

**Extended Data Figure 4.**
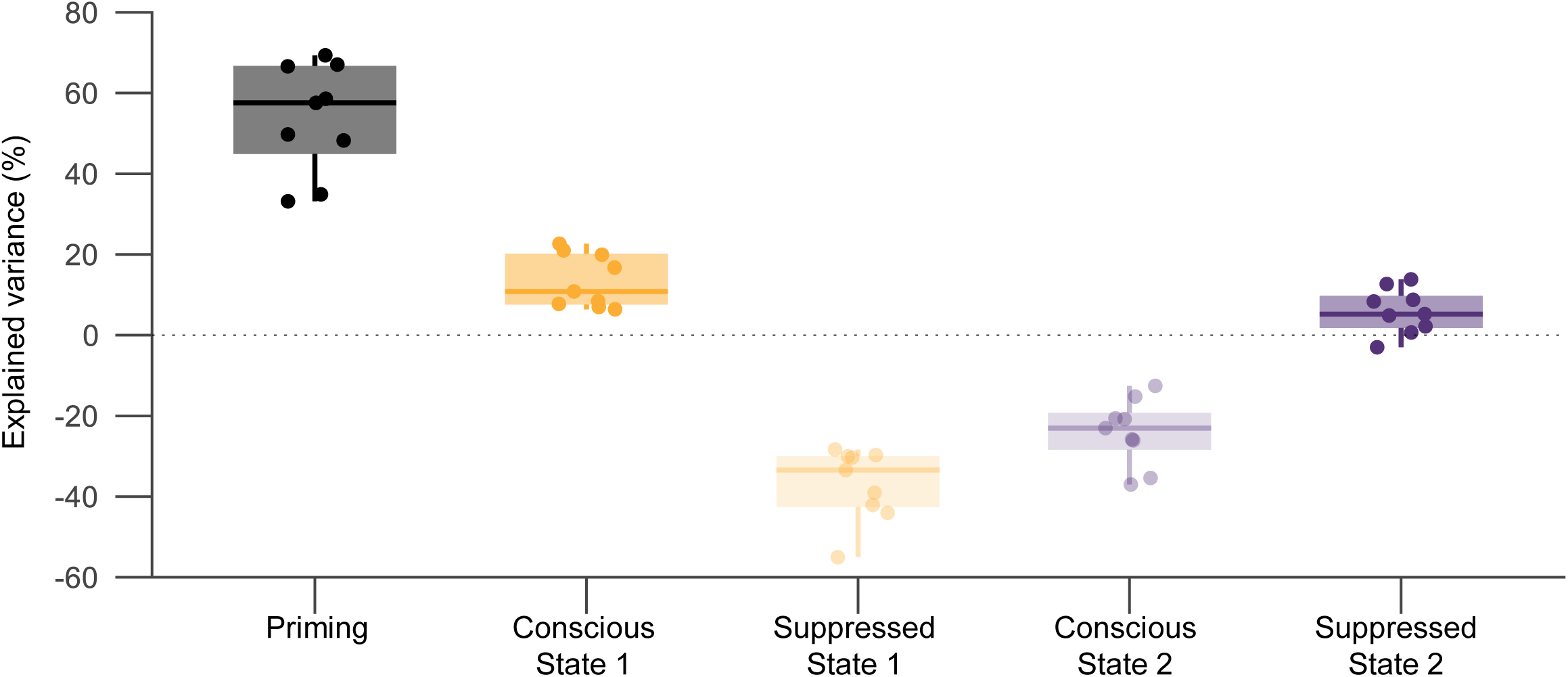
Interleaved representations in individual sessions. Explained variance (mean of the shape and appearance explained variances, each image-variance-weighted across its features) for nine individual sessions. Boxes, median and interquartile range across the 9 sessions; whiskers, range; points, individual sessions. As in Fig. 3, states were inferred from one half of neurons and explained variance was computed using the held-out half of neurons.

**Extended Data Figure 5.**
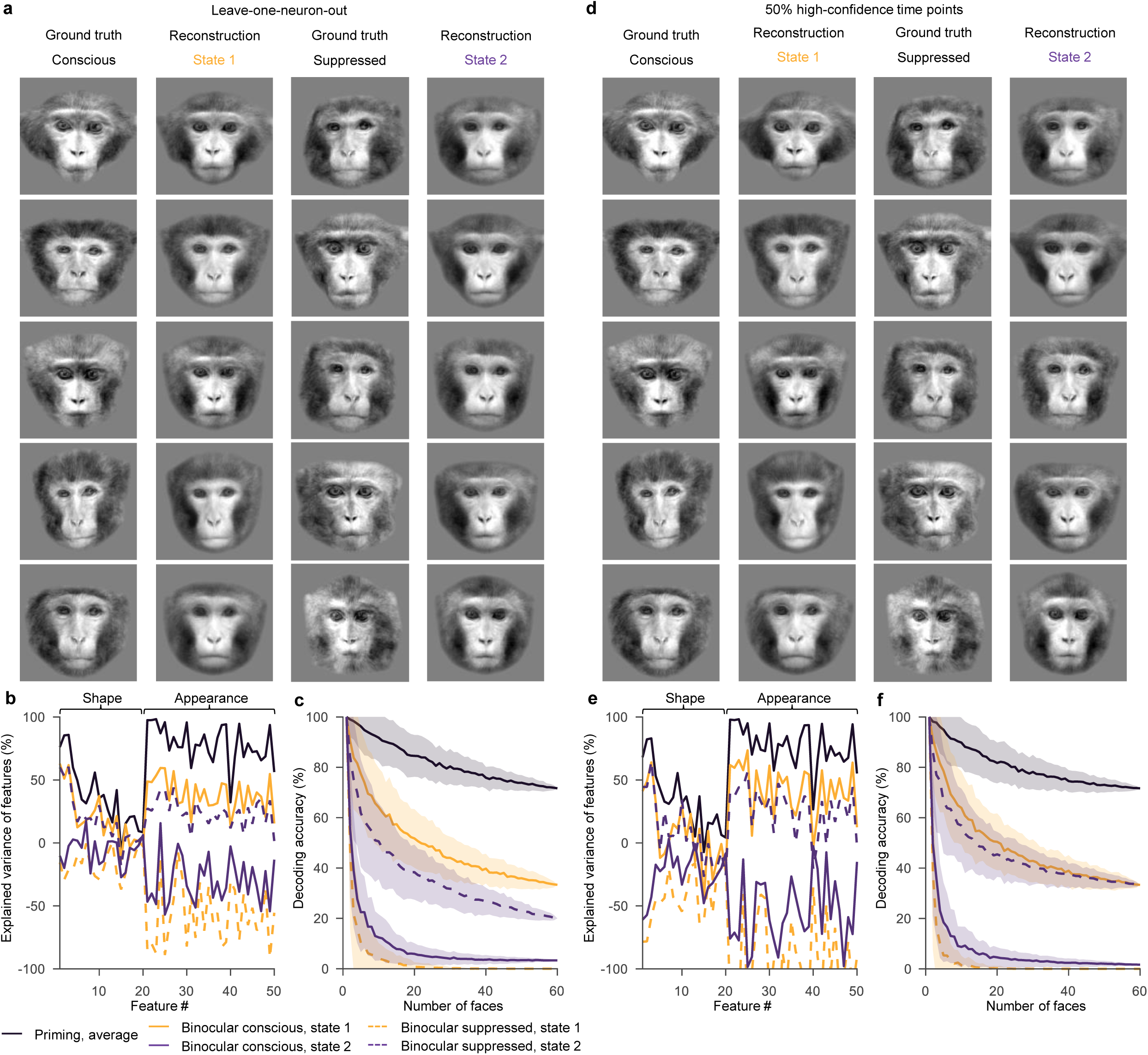
Interleaved representations using leave-one-neuron-out cross-validation and high-confidence time points. (**a**)-(**c**) Leave-one-neuron-out cross-validation. Same conventions as Fig. 3e-g but for each neuron the state inference was computed using all other neurons, and then responses of each neuron were pooled to reconstruct faces. (**d**)-(**f**) Reconstructions from high-confidence time points. Same conventions as Fig. 3e–g, but instead of assigning all time points to either state 1 or state 2, for each state the analysis was restricted to the 50% of time points with the largest absolute-value log-likelihood ratio (i.e., the most confidently assigned time points).

**Extended Data Figure 6.**
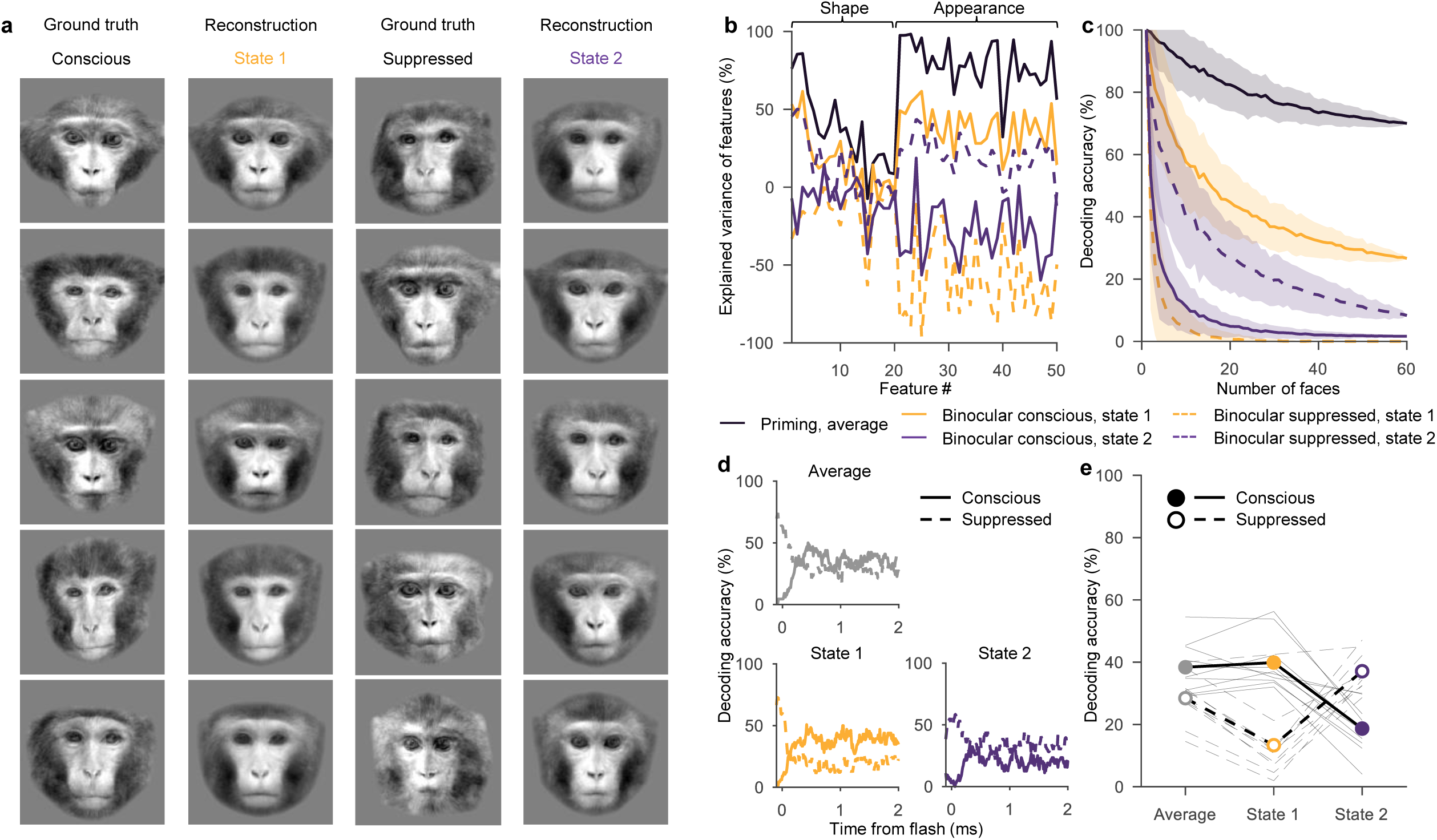
Balance-trial control. (**a**)-(**e**) Same conventions as Fig. 3e-i but for subsampling time points so that for each trial there is the same number of state 1 and state 2 time points. As in Fig. 3, states were inferred from one half of neurons and reconstruction and decoding analyses performed on the held-out half.

**Extended Data Figure 7.**
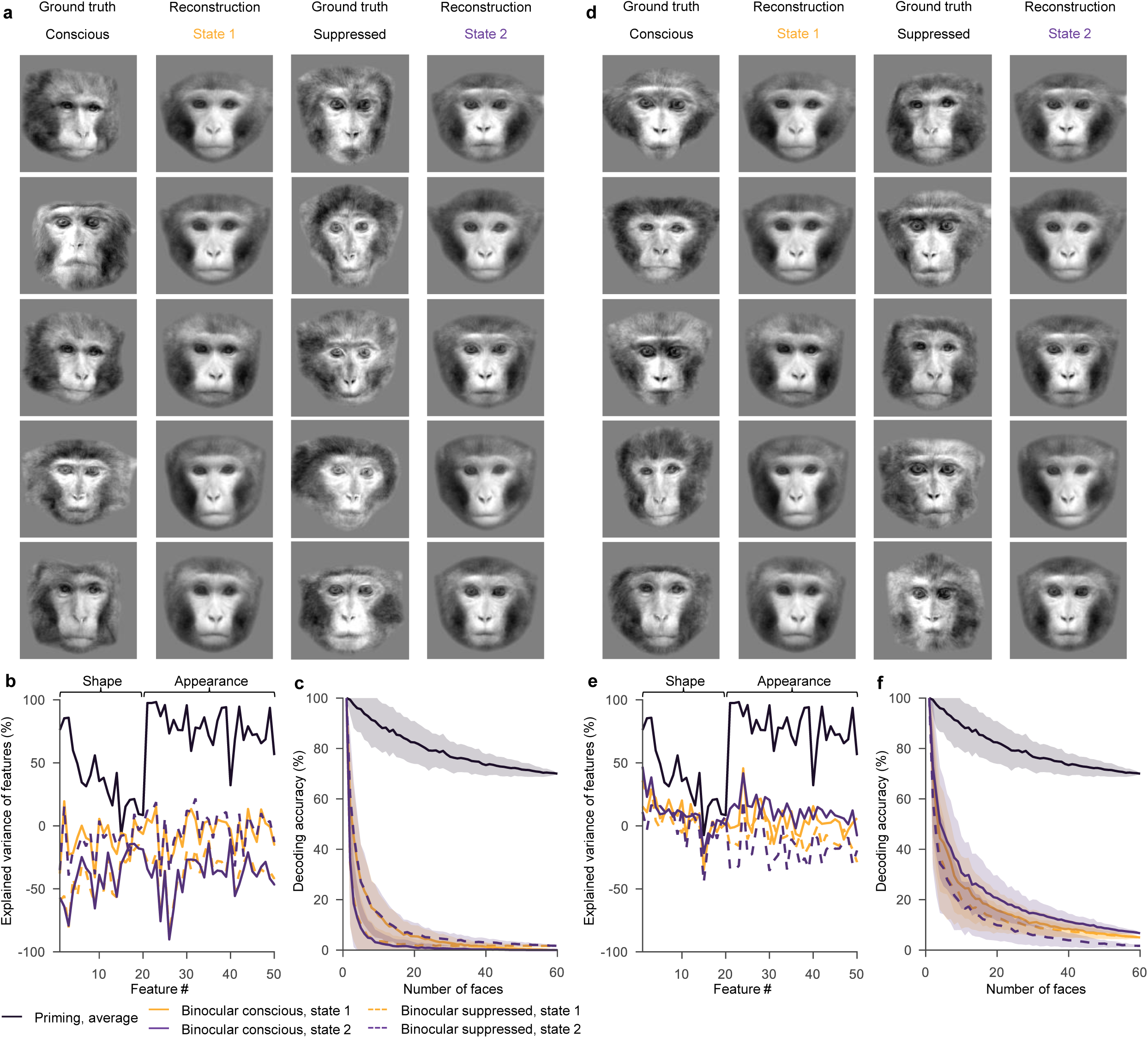
Shuffle template control. (**a**)-(**c**): Same conventions as Fig. 2h-j but response templates for inferring states were randomly reassigned. For each presented stimulus pair (A, B), the templates were replaced by those of two randomly selected stimuli (A′, B′). The same stimulus reassignment was used across neurons, trials and sessions. Reconstructions were then compared to these newly assigned templates (A’, B’). (**d**)-(**f**) Same as (a)-(c) but reconstructions were compared to the actually presented faces (A, B). Reconstructions for this control were obtained using circular reuse of the data for state estimation (double-dipping, as in Fig. 2) rather than cross-validation, so as to yield the highest-quality reconstructions and thus the most stringent control.

**Extended Data Figure 8.**
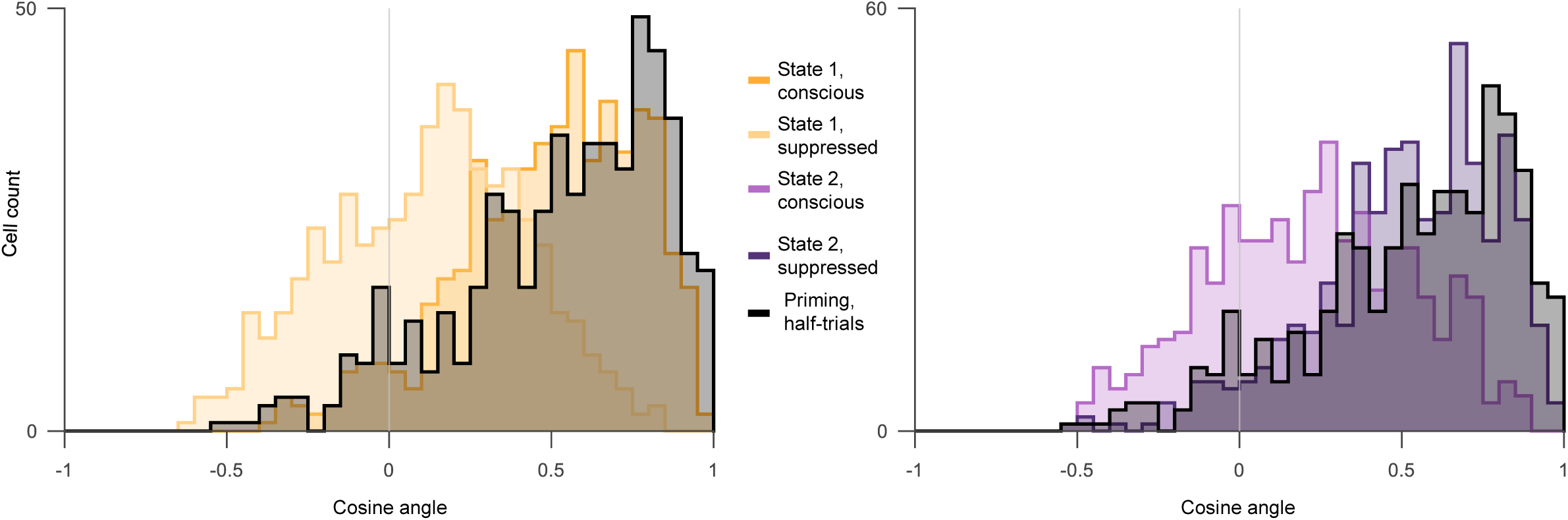
Single-cell feature-tuning axes within each state align with the priming axis. For each cell with priming-reconstruction explained variance > 0.05 the feature-tuning axis was estimated either from its priming responses (as in Fig. 4) or from its state-resolved binocular responses to either the conscious or suppressed face. Alignment between a state axis and the priming axis is quantified as the cosine similarity between the two weight vectors (intercept excluded); histograms show its distribution across 548 selected cells. Left, state 1; right, state 2. Within each panel, the axes estimated when the cell’s preferred face was conscious and when it was suppressed are shown separately with different colours, see legend. We also computed the noise ceiling of the priming tuning axis itself, as the cosine similarity between priming axes estimated from two independent halves of the trials (black distribution). The axis computed from the conscious face during state 1 was only slightly less aligned with the priming axis than the noise ceiling (two-sided paired t-test, *t*(547) = −2.5, *p* = 0.01, *n* = 548 cells). The same was true for the state 2 suppressed face (two-sided paired t-test, *t*(547) = −0.9, *p* = 0.35, *n* = 548 cells). In comparison, the alignment was much weaker for state 1 suppressed compared to priming noise ceiling (two-sided paired t-test, *t*(547) = −24.0, *p* = 10^−87^, *n* = 548 cells) and for state 2 conscious (two-sided paired t-test, *t*(547) = −20.3, *p* = 6 × 10^−69^, *n* = 548 cells). The first principal component of the face features, corresponding to stimulus colour, was removed, as in Fig. 4a-f.

**Extended Data Figure 9.**
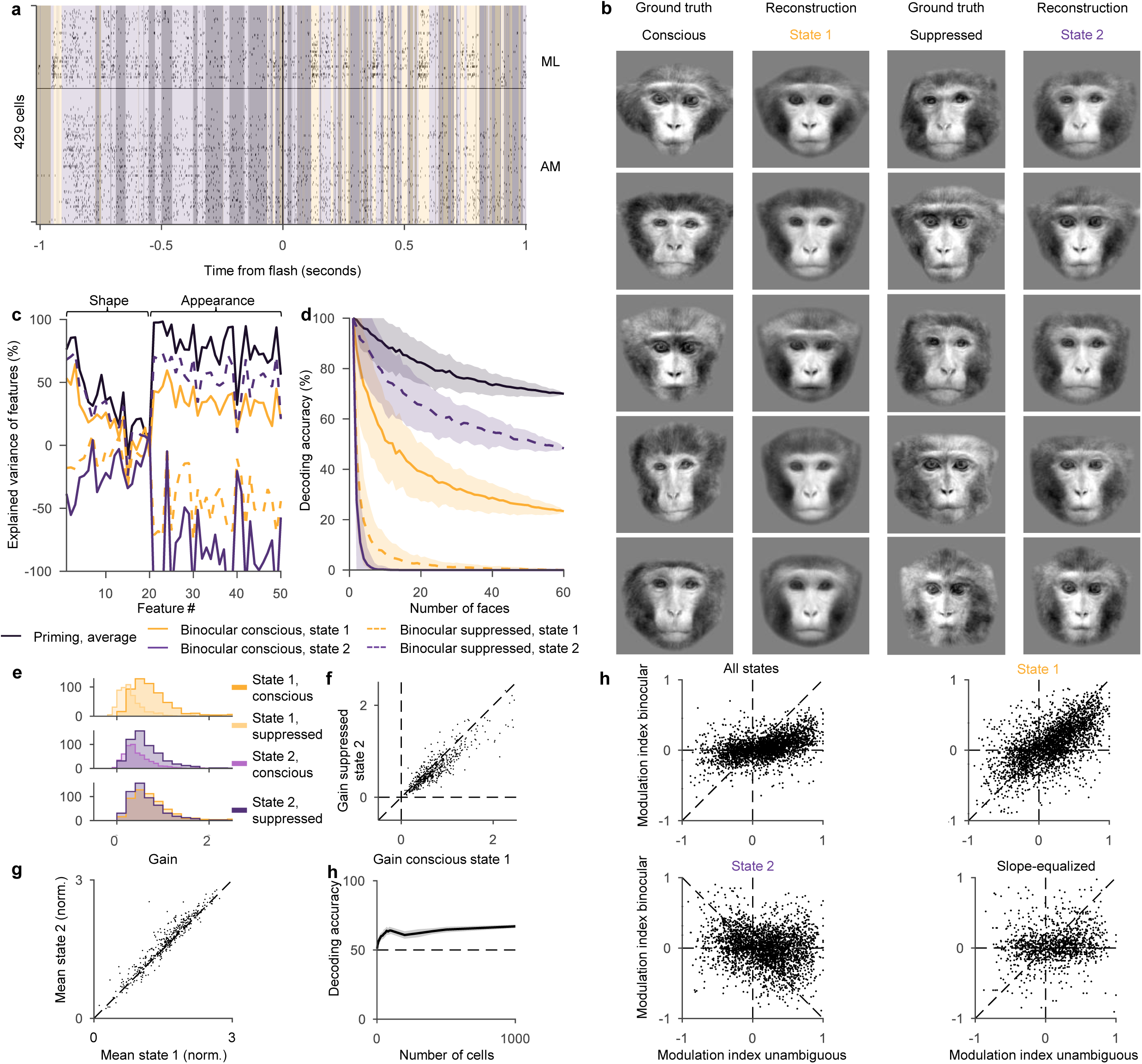
On/Off states are separate from conscious/suppressed states. (**a**) Example raster as in Fig. 1g. Orange and purple overlays correspond to conscious and suppressed states. Bright and dark overlays correspond to On and Off states. (**b**)-(**d**) Same as Fig. 3e-g but for a four-state LLR that included On/Off x conscious/suppressed states. State 1 = conscious On+Off states, State 2 = suppressed On+Off states. As in Fig. 3, states were inferred from one half of neurons and reconstructions obtained using the held-out half. (**e**)-(**h**) Same as Fig. 4c-g but for four-state LLR.

**Extended Data Figure 10.**
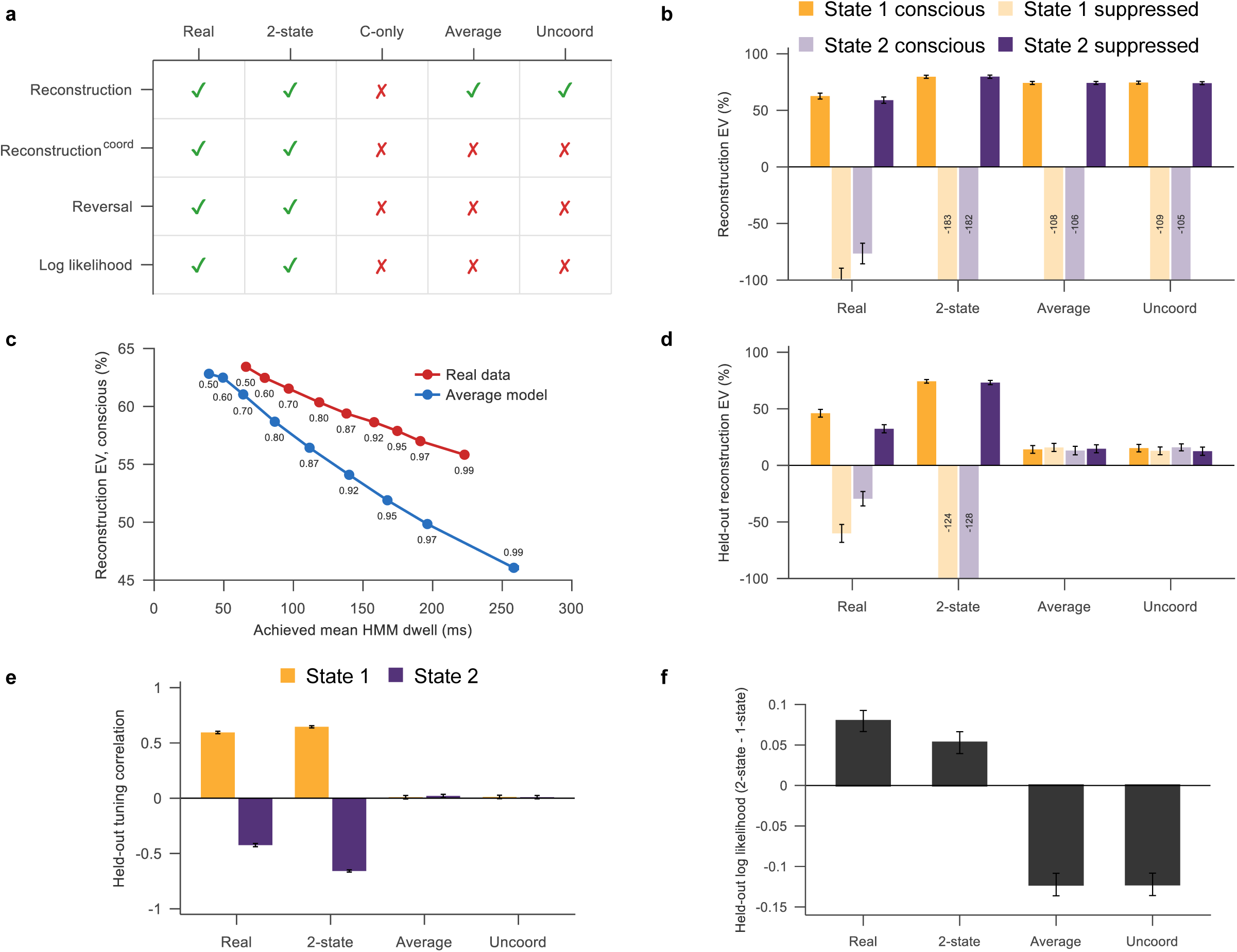
Comparison of the two-state model to alternative models. Across all panels, models are abbreviated: Real, recorded data; 2-state, two-state; C-only, conscious-only; Average, averaging; Uncoord, uncoordinated. (**a)** Model × test matrix. Columns, the recorded data and the four generative models; rows, the four tests. Green check, pass; red cross, fail. Pass criteria: Reconstruction/Reconstruction^coord^, in state 1, the explained variance of the conscious face exceeds both zero and that of the suppressed face; in state 2, the explained variance of the suppressed face exceeds both zero and that of the conscious face (bootstrap for each comparison, *p* < 0.05). The superscript “coord” denotes the cross-validated version of reconstruction, in which the state is inferred from one half of the neurons and reconstructions are evaluated in the held-out half. Reversal, state 2 held-out tuning correlation significantly negative (bootstrap, *p* < 0.05). Log-likelihood, held-out two-state log-likelihood exceeding one-state. (**b**) Reconstruction accuracy. Explained variance for the conscious and suppressed face features, reconstructed from state-resolved activity (mean of the shape and appearance explained variances, each image-variance-weighted across its features), for the recorded data and each model. Bars, mean; error bars, bootstrap standard deviation over stimulus pairs (2000 samples). (**c**) Conscious-face reconstruction explained variance in state 1 as a function of the achieved mean state dwell time, for the recorded data (red) and the averaging model (blue). Each point corresponds to one imposed HMM self-transition probability *p*; *p* was varied from 0.5 to 0.99 and for each *p* the empirical dwell time was measured for both real and synthetic data (x-axis). (**d**) Cross-validated reconstruction accuracy. As in b, but with the state inferred from one half of the neurons and the face features reconstructed from the held-out half. (**e**) Held-out tuning reversal. State-1 (orange) and state-2 (purple) single-cell tuning correlation *r* for the recorded data and each model. States were inferred from one half of the neurons and tuning measured in the held-out half. Bars, mean; error bars, standard deviation of 2000 bootstrap resamples across cell-stimulus pairs. (**f**) Model comparison. Difference in cross-neuron validated conditional log-likelihood between a two-state and a one-state Hidden Markov model (per cell) for the recorded data and each model; values above zero indicate that two states fit better. Error bars, s.e.m. across sessions.

## Methods

All animal procedures in this study complied with local and National Institutes of Health guidelines including the US National Institutes of Health Guide for the Care and Use of Laboratory Animals and were performed with the approval of the California Institute of Technology Institutional Animal Care and Use Committee (IACUC) and the University of California Berkeley Animal Care and Use Committee (ACUC). Human participant experiments were performed with approval of the California Institute of Technology Institutional Review Board (IRB).

### Targeting

Two male rhesus macaques were implanted with head posts and acrylic implants. We targeted face patches ML and AM in IT cortex, as well as ventrolateral prefrontal and orbitofrontal cortex for electrophysiological recordings. ML and AM were identified using fMRI. Monkeys were scanned in a 3T scanner (Siemens), as described previously^31^. During functional scans, MION contrast agent was injected to increase signal-to-noise ratio and monkeys passively viewed blocks of faces and blocks of other objects to identify face-selective patches in the brain, using a custom eight-channel coil (MGH) at isotropic 1 mm resolution. Recording chambers were implanted over ML and AM, and custom grids and lattices were 3D printed (3D Systems) and inserted into the chamber to target the desired areas. Both chamber placement and grid design were planned with the software Planner^32^. We inserted tungsten electrodes and then performed anatomical MRI scans to confirm correct targeting of the desired location.

### Electrophysiological recording

Single tungsten electrodes were used for scout recordings in the targeted areas prior to Neuropixels recordings. An Omniplex DHP system (Plexon) and custom analysis software written in MATLAB (MathWorks) were used for these initial recordings. Tungsten electrodes with an impedance of 100 kΩ (FHC) were inserted through plastic guide tubes (WPI), that extended 2 mm past the dura. Guide tubes were inserted through a custom 3D printed grid with holes designed to be in the direction of the target. After tungsten electrode recordings confirmed the presence of cells at the desired target, custom 3D-printed lattices were designed for Neuropixels recordings using the same trajectory. Subsequently, Neuropixels 1.0 NHP probes (45 mm long probes with 4416 contacts along the shaft, of which 384 are selectable at any time) were inserted using a custom-machined insertion mechanism, as part of which either a MDO-974A microdrive (Narishige) or an ultrasonic piezo motor drive (XLA-1 series, Xeryon) slowly advanced the Neuropixels probe through a stainless steel cannula (A-M systems) along a linear ball bearing rolling guide rail (IKO). Our insertion mechanism is described in more detail in ^13^. All Neuropixels data were acquired using SpikeGLX (Bill Karsh, HHMI/Janelia Research Campus) and spike sorted using Kilosort3^33^. Both single-units and multi-units were used for analyses.

### Stimuli and behavioural task

Monkeys were head-fixed and viewed an LCD screen (Asus ROG Swift PG43UQ) of 60 visual degree width and 44 degree height in a dark room. Stimuli were displayed on the screen with a 7 visual degree size while red/cyan anaglyph goggles were placed in front of the left and right eye of the monkey. Anaglyph goggles were custom made with filters (LEE filters) to match the red and green/blue emission spectrum of the screen, respectively, so that inputs to left and right eyes could be controlled independently. Emission spectra were measured using a PR-650 SpectraScan colorimeter (Photo Research). Gaze position was monitored using an infrared camera eye tracking system (Eyelink) sampled at 1000 Hz. The camera recorded one or both eyes through the red/cyan anaglyph filter.

In the first phase of the experiment, each monkey passively viewed two sets of 96 and 49 screening stimuli, in pseudorandom order (150 ms ON time, 150 ms OFF time) with a fixation spot of 0.2° diameter in the center of the screen. Screening stimuli consisted of 16 images of faces and 80 non-face images in the first screening set, with ∼8 repeats per image, and 40 images of faces and 9 non-face images in the second screening set, with ∼30 repeats per image. Monkeys received a juice reward for maintaining fixation for at least 3 s within a 0.5° rectangle centered on the fixation spot. Stimulus progression was contingent on fixation. For all experiments, only trials where monkeys maintained fixation for at least 90% of the trial duration were included for analyses.

After the screening phase, one of two types of flash suppression stimulus sets was presented. In flash suppression, a priming stimulus is presented, followed immediately, without a blank period, by a binocular stimulus or another type of control stimulus (**Fig. 1a**). For the first type of stimulus set, ten face/object stimuli (five stimuli for left eye, and five stimuli for right eye) were selected from the 49 stimuli screening set that maximized the mean pairwise Euclidean distance between stimuli in neural space. The 25 combinations of the five left eye stimuli and five right eye stimuli were presented as stimulus pairs. For each stimulus pair, one of the two stimuli was presented monocularly as a priming stimulus for two seconds. The priming stimulus was followed by one of the two types of test stimuli, presented for four seconds: (1) a binocular stimulus, consisting of the priming stimulus and the other stimulus of the stimulus pair presented to the two eyes, respectively, referred to as the ‘binocular’ context, (2) a monocular stimulus, consisting of only the other stimulus of the pair that was not presented during priming, referred to as the ‘unambiguous’ context, used for the modulation on the x-axis of **Fig. 1h** and **Fig. 4g**. For each stimulus pair, both contexts were presented, and either stimulus was presented as priming stimulus in different trials. The new stimulus that was flashed during the post-priming test stimulus is referred to as the dominant stimulus, whereas the stimulus that was already present during priming is referred to as the suppressed stimulus. The test stimulus was followed by a 500 ms blank period containing only the fixation spot on a grey screen. Four stimulus pairs were presented with ∼30 trials per condition, and the rest of the 25 stimulus pairs were presented with ∼5 trials per condition (for image pair × which image was dominant × unambiguous / binocular = 100 conditions). This experiment was performed in seven sessions in monkey A and three sessions in monkey M and analysed for **Fig. 1h**, **Fig. 2c,d,k,l, Fig. 3a,h,i, Fig. 4g**, **Fig. 5c,d** and **Fig. 6f**.

For the second type of flash suppression stimulus set, 1500 unique faces, randomly sampled from face feature space, were presented. The same 1500 faces were presented in all experiments using this second type of stimulus. To maximize the number of unique stimuli, we did not present all combinations; instead, this set consisted of 750 stimulus pairs, with each of the 1500 faces being part of only one stimulus pair. Moreover, the priming stimulus duration and post-priming test stimulus duration were both reduced to one second, and the blank period after the test stimulus was reduced to 250 ms. As in the first experiment, both stimuli of the stimulus pair were presented as priming stimulus in different trials. This type of experiment did not include an unambiguous alternation condition. A subset of 60 faces was presented for a higher number of repeats (∼20 trials per condition for high repeat stimulus pairs, ∼2 trials per condition for the rest of the 750 stimulus pairs). Reconstructions were generated for this subset of 60 faces, while the linear regression was trained on all 1500 faces. This experiment was performed in a separate set of five sessions in monkey A and four sessions in monkey M, and analysed for **Fig. 1g**, **Fig. 2b-j**, **Fig. 3a-g**, **Fig. 5** and **Fig. 6a-e**.

### Data analysis

For fitting LLR and HMM, Python version 3.11 was used. All other analyses were performed using MATLAB 2023b and 2025b (MathWorks). For **Extended Data Fig. 5** and **Extended Data Fig. 10** Claude Code was used in order to reduce computational runtime, and all generated code was reviewed for correctness by J.K.H.

#### Human reports

Because button reports lag stimulus presentation, we first estimated report latency from the priming period, independently of the flash: for each observer, the time at which the probability of reporting the primed colour reached half its rise from the pre-onset baseline to its maximum (0.86 ± 0.05 s, mean latency ± s.e.m., **Fig. 1b**). The analysis window for determining significance of suppression was 300–1000 ms after flash onset (the same window as used for neural analysis, see below), shifted by this latency. Flash suppression was quantified per observer as a suppression index: the mean difference in report probability between the flashed and the priming stimulus within this window. We performed a two-sided t-test of this suppression index against the null hypothesis of it being 0 across all 14 human participants. The flashed percept was reported on 82% of binocular rivalry trials, only 5 percentage points below the 87% correct reporting of the unambiguous priming stimulus. The 13-percentage-point deviation from ceiling during priming provides an estimate of baseline report errors human participants made even when the percept was unambiguous. The small additional reduction during rivalry may be partially due to a refractory period in reporting rather than failures of perceptual suppression: rivalry reports followed shortly after the preceding priming report, whereas the priming report was made after a substantially longer response-free interval. Closely spaced responses are known to interfere with one another, producing a psychological refractory period or dual-task cost that reduces accuracy of the second response^34^.

#### Log-likelihood ratio (LLR)

For each binocular trial we computed a continuous single-trial index of the momentary state as the population log-likelihood ratio (LLR) between the two states of a Poisson mixture model with equal mixture weights (**Fig. 2a**). In consecutive 10-ms bins, the spike counts of the visually responsive IT population were evaluated under the two emission templates, which were computed from the mean firing rates of each neuron during the corresponding single-face priming trials. The LLR was computed as the difference of the population Poisson log-likelihoods, 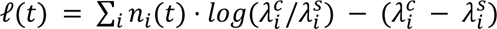, where *n_i_*(*t*) is the spike count of neuron *i*, 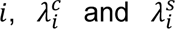 its template firing rates derived from priming activity when the conscious/suppressed face was presented unambiguously, and *Δt* = 10 *ms*. l(*t*) is positive when the population evidence favours the conscious state and negative when it favours the suppressed state, and states were assigned based on this sign.

To determine whether rivalry responses were better described by two states than by a single static response, we compared the marginal likelihood under the two-state Poisson mixture model (used for LLR decoding) with that under the best-fitting one-state model. The best-fitting one-state model was obtained by maximum-likelihood estimation, yielding a static firing-rate vector equal to the average response during binocular rivalry trials (**Fig. 3a**). For the two-state model, the two states were assigned equal prior probability and the marginal likelihood of the spike-count vector in each time bin was computed by averaging its Poisson likelihood under each of the two priming templates, thereby marginalizing over state assignments. The likelihood of a trial was the product of these marginal likelihoods across time bins. To compute the likelihood of the static model, we split the rivalry trials into halves, computed the average rivalry response from one half, and then calculated the Poisson likelihood of the spikes in the other half using this estimate as the underlying firing-rate vector. We then repeated this procedure with the two halves reversed. The two-state templates were estimated from independent priming trials and therefore did not require a trial split.

#### Autocorrelation and LLR timescale

For each session, we computed the autocorrelation of the LLR at 1 ms lag resolution up to 500 ms, pooling over binocular trials. We then averaged this curve across 19 sessions (**Fig. 2d**). As a null, we also recomputed the autocorrelation of the LLR after permuting time bins of neural data within trials, independently for each neuron, which abolished all temporal structure. Both real and shuffled autocorrelation curves were normalized by their zero-lag value. The real autocorrelation has two regimes: below 10 ms it reflects the triangular autocorrelation of the 10-ms analysis window (l is constant within a bin) and carries no dynamical information, whereas beyond 10 ms it decays with the persistence of the LLR signal. The shuffled autocorrelation goes to zero after the 10 ms window, as expected. Modelling the state as a symmetric two-state Markov chain, whose autocorrelation is *ρ* · *e*^−2*τ*/*D*^ with mean dwell time *D*, we fit this form to the session-averaged autocorrelation over lags 10–90 ms (from the bin edge to the first zero-crossing) by least squares, with the amplitude ρ free. This mean dwell time of *D* = 67 *ms* corresponds to a median dwell of *D* · *ln*2 ≈ 46 *ms* for exponentially distributed dwell times, approximately matching the HMM-inferred time scale described below.

#### Reconstructions

Face features were predicted from population activity by linear regression, *F* = *W* × *R*, where *F* is the vector of 50 face features, *R* is the population response projected into principal component (PC) space, and *W* is a matrix of regression weights. Responses of 3501 visually responsive neurons across sessions were computed from spike counts averaged across trials and across time points from 300 ms to 1000 ms after stimulus onset and baseline-normalized using the spike counts from 50 ms after blank onset until the end of the blank period that followed each binocular flash suppression stimulus. Visual responsiveness of cells was independently determined based on whether responses from 70 to 200 ms after stimulus onset were significantly different from baseline firing 0 ms to 70 ms after stimulus onset during the first stimulus set with 96 stimuli (t-test, *p* < 0.05).

Because the number of neurons greatly exceeded the number of stimuli available for fitting, regressing features directly onto single-neuron responses would have overfitted the data. To prevent overfitting, we first projected neural responses into a 500-dimensional space using principal component analysis (PCA) before performing linear regression (MATLAB lscov, ordinary least squares, no regularization); 300 components were used when reconstructing from the 661 prefrontal neurons, reflecting the smaller population. Coefficients for PCA were computed from priming responses, and subsequently these coefficients were used to project all responses into the same space. Linear regression weights were also computed using priming responses, and then those same weights were used when predicting face features in the binocular context. During the binocular context, only time points during the same time window 300 to 1000 ms that were assigned to state 1, or state 2, respectively, were used to compute responses, whereas for priming responses all time points from 300 ms to 1000 ms were averaged. For both priming and state-resolved binocular rivalry, predictions were cross-validated by excluding the predicted face from the training set. After averaging state-dependent responses for each neuron, responses were concatenated across the neuron dimension across sessions. Reconstructions of face images were computed from face features using the active appearance model used in ^35^, rendered in greyscale and contrast-normalized for visualization purposes (**Fig. 2h**).

#### Explained variance

For the explained variance of a given feature, we computed one minus the ratio of the sum of squared residuals and the sum of squared mean-subtracted features (**Fig. 2i**): 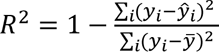, where *y_i_* is the actual feature of face i, 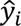 is the predicted feature of face i, and *y* is the mean of the feature across faces. To test whether the two states encode the conscious and suppressed faces separately, we reconstructed the face features in each state and, for each stimulus, computed a leaning index: the distance (in feature space) from the state’s reconstruction to the suppressed face minus its distance to the conscious face (positive = closer to the conscious face; conscious = reported/dominant face, suppressed = its pair partner). We then took the difference between states, leaning(state 1) − leaning(state 2), which is positive when state 1 leans toward the conscious face and state 2 toward the suppressed face. This difference was averaged over each pair’s two dominance conditions and tested against zero with a one-sample t-test across high-repeat stimulus pairs. To aggregate the explained variance of individual face-model features into a single metric, we computed the explained variance separately for the shape features and the appearance features and averaged the two. Within each set, features were weighted by the variance they contribute to the face image, namely the variance in landmark coordinates for shape, and in the standard-shape pixel intensities for appearance.

#### Euclidean decoding

Euclidean-distance-based decoding was performed on the predicted face features (**Fig. 2j**). We picked the 50 principal z-scored shape and appearance features that were predicted for a given face, and then compared the Euclidean distance of the selected features to the ground truth features of the selected face and the ground truth features of a given number of randomly chosen distractor faces. The selected face was labelled as decoded correctly if the distance of the predicted features was closer to its ground truth features than features of all the other faces. This was repeated for all 60 predicted faces to yield a decoding accuracy.

To test for statistical significance of the Euclidean-distance-based decoding we computed for each high-repeat face identity the 10-way Euclidean nearest-neighbour decoding accuracy of the state-resolved reconstruction against the true face-feature vectors (averaged over random 10-identity subsets), separately for state-1 (dominant/conscious) and state-2 (suppressed) responses and for the dominant and the suppressed face. Per identity we formed the interaction contrast, (dominant_state 1_ –suppressed_state 1_) – (dominant_state 2_ − suppressed_state 2_), averaged the two members of each rivalry pair into a single value (n = 30 pairs, the unit of inference), and tested it against zero with a two-sided sign-flip permutation test (10⁵ permutations, each randomly flipping the sign of whole pairs).

#### Single-trial decoding

A cross-validated multi-class classifier was trained to read out face identity from the population response and applied to individual binocular trials. The classifier (a linear support-vector machine, 8-fold stratified cross-validation) was trained on the priming trials, using as features each neuron’s mean firing rate over the priming response window; class labels were the identities of the 10 presented faces.

The priming-trained classifier was then applied, without retraining, to binocular trials, in sliding 200 ms windows stepped by 10 ms across the trial (**Fig. 2k**) or over the full stimulus window (**Fig. 2l**). For every trial and window, the population response was averaged over the window and classified into a single predicted face. We computed conscious decoding accuracy, the fraction of trials on which the predicted face matched the consciously perceived face, and suppressed decoding accuracy, the fraction matching the suppressed face.

We repeated the readout on state-restricted responses: within each window the population response was averaged only over time points assigned to a given state (state 1 or state 2), and trials with no time points in that state in the window were dropped.

#### Axis tuning

For each neuron we defined its preferred axis in face-feature space from a cross-validated (leave-one-face-out) linear encoding model fit to average priming responses, predicting the neuron’s response to each face from face features; the fitted weight vector is the neuron’s preferred axis, and the predicted response is each face’s projection onto it.

For the example cell (**Fig. 4a**), tuning was visualized in two feature dimensions: the preferred axis (x, cross-validated predicted response) and an orthogonal axis (y, first principal component of the features after orthogonalizing to the preferred axis; the PCA was computed on stimuli with the same colour so that it did not collapse onto the colour axis). Each point corresponds to one face image, positioned by its projection onto the two axes and coloured by the neuron’s measured response (red: high response, blue: low response) during the binocular condition in a given state/percept; accompanying marginal plots show the neuron’s binned response along the preferred axis for state 1/conscious (solid) versus state 2/suppressed (dashed).

For population tuning maps (**Fig. 4b**), each neuron’s state-resolved binocular response was plotted across its axis-ranked stimuli as a neurons × stimuli image, separately for the four conditions — state 1 / conscious face, state 2 / conscious face, state 1 / suppressed face, state 2 / suppressed face. We only included neurons whose priming responses could be explained by features (cross-validated *R*² > 0.05). Axis tuning gain was quantified per neuron as the slope of an ordinary-least-squares regression of the measured response on stimulus position along the preferred axis (intercept + slope), computed separately in each state and percept.

#### Decoding state 1 vs state 2

To test whether IT population activity allows distinguishing the two inferred states, generalizable beyond stimulus-specific tuning, we trained a linear decoder to discriminate the ordering of paired state 1 and state 2 responses (**Fig. 4f**). For each stimulus we took the population response vectors to that stimulus being conscious during state 1 and to that stimulus being suppressed during state 2. We then constructed a balanced two-class decoding problem: for every stimulus, we created two types of population vector concatenations: the state-1 response, followed by the state-2 responses, was labelled class 1, and the reversed concatenation, state 2, followed by state 1, was labelled class 2. Because each stimulus contributes identical responses to both classes and differs only in concatenation order, stimulus identity carries no information about the label; the only cue is whether the state-1 response is the first half or second half of the vector. The decoder can therefore only discriminate labels correctly if the state-1 and state-2 responses to the same stimulus are systematically distinguishable, beyond stimulus identity. We trained a linear SVM on a random half of the stimulus pairs and tested on the held-out half with both members of a pair kept in the same set, so that no stimulus appeared in both training and test. Accuracy was computed as a function of number of neurons by subsampling neurons, over 100 random split-and-subsample iterations per size.

#### Modulation

Modulation indices were computed using the average response for each cell and stimulus pair from 300 ms to 1000 ms after stimulus onset, where human participants reported only perceiving the flashed stimulus for most trials after accounting for report delay. We chose this time window conservatively: 300 ms avoids visual transients caused by the flash, as visual response latencies are around 100 ms in face patches. The end of the time window, 1000 ms, was determined by the stimulus duration in the main experiment. We computed modulation indices using the formula 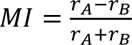, where *r* and *r* are the responses to stimulus A and B, respectively. For the x-axis, the responses to unambiguous stimuli (A, B) of a given pair were used, while for the y-axis, the responses to binocular stimuli were used, when stimulus A was dominant for *r_A_* and when stimulus B was dominant for *r_B_*. For **Fig. 4g**, the responses for modulation indices on the y-axis were computed over time points from 300 ms to 1000 ms that were assigned to state 1 or state 2, respectively, whereas modulation indices on the x-axis were computed across all timepoints. We inferred states using one half of neurons and computed modulation indices only for the held-out set of neurons. Only cells that (1) were significantly selective to A vs B during priming responses (t-test over responses 300 ms to 1000 ms after priming onset, with threshold *p* < 0.05) and (2) had at least one spike for both stimuli of the pair and both states were included. These selection criteria included a total of 2885 cell-stimulus pairs. For gain-equalization, we computed tuning gain for state 1 conscious and state 2 suppressed, and modified responses during state 1 to equalize the gain: 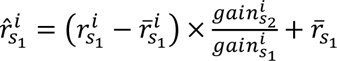, where 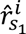 is the gain-normalized response of neuron *i* to the conscious face in state 1, 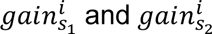 are the tuning gain of neuron *i* to the conscious face in state 1 and suppressed face in state 2, respectively, and 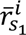 is the mean response of neuron *i* during state 1, which is kept constant during the gain equalization. Neurons whose two gains had opposite sign, for which no positive rescaling exists, were excluded from this control. The gain-equalized average response to binocular rivalry was then computed as 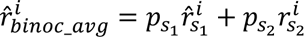 where 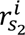 is the non-gain-equalized response to the suppressed face during state 2, and *p_s_*_1_ and *p_s_*_2_ are the proportions of inferred states 1 and 2 for the given stimulus pair. The relationship between binocular and unambiguous face selectivity was quantified as the slope of each neuron’s binocular modulation index against its unambiguous modulation index, estimated by total least-squares regression. Because both modulation indices are noisy single-neuron estimates, we used total least squares, which is symmetric in the two variables rather than ordinary least squares which would be attenuated toward zero by measurement noise.

#### Hidden Markov Model (HMM)

HMMs with two latent states were fit to neural recordings to partition each trial into segments representing the conscious and suppressed stimulus, respectively (**Fig. 5a-d**). For each stimulus pair (A, B), average stimulus response vectors *r^A^* and *r^B^* were computed from priming trials so that *r_i_^A^* is equal to the average firing rate of unit *i* across all priming trials where stimulus A was shown. To account for stimulus onset time, only time bins 70-1000 ms after stimulus onset were included in this average. These average response vectors were then used as fixed firing rates in a two-state Poisson HMM for identifying micro-epochs in this stimulus pair. The hidden state *z_t_* ∈ {1, 2} for this model follows a Markov chain with transition probabilities determined by a 2 × 2 matrix *T*:

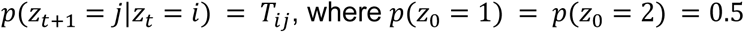

The vector of unit spike counts *y_t_* ∈ *R^N^*for time *t* with a bin size of *Δt* is assumed to follow a Poisson distribution for each unit with mean determined by the state *z_t_*:

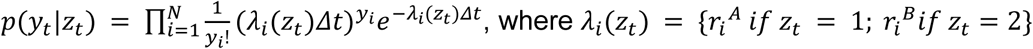

The transition matrix T is the only free parameter in this HMM, and was fit by maximizing the likelihood of binned spike counts observed during binocular trials for each stimulus pair, using the Baum-Welch algorithm. After the transition matrix was estimated, the Viterbi algorithm was used to compute the most likely state sequence {*z_k_*_1_, *z_k_*_2_, … *z_kT_*} for each binocular trial *k* for all stimulus pairs. The HMM states {*z_kt_*} corresponding to the stimuli were converted into states {*c_kt_*} indicating whether the dominant or suppressed stimulus was being represented at time *t*:

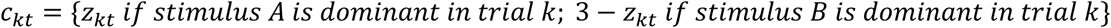

To compute conditional likelihood, we split units into two random halves. We used the first half to infer states with the HMM, and then calculated the likelihood of observing the spikes from the second half of units given the state assignments inferred from the first half. To compare to a single-state model, we then calculated the likelihood of observing the same spikes from the second half under a static rate, namely, the average of the priming templates. For each session, we took the trial-averaged difference between the HMM model and the static model and normalized it by the number of units used to compute it.

All code for HMM fitting and prediction was written in Python using the hmmlearn package.

#### Synthetic models

We simulated four generative models of the single-trial population activity, each constructed per session from the priming-derived response templates of the two faces of a trial (*r_conscious_*, *r*_suppressed_) and matched to the recorded number of trials and time bins of each stimulus pair (**Extended Data Fig. 10**). In every model, spike counts in each bin were independent Poisson draws based on the mean firing rate for that bin. The only difference between models was how that mean firing rate was set for each time bin and each neuron. Two-state (temporal multiplexing): the entire population followed a single hidden two-state Markov chain with a self-transition probability giving a mean dwell of 60 ms, held identical across sessions, emitting from the conscious template in one state and the suppressed template in the other. The same time partition was used for all neurons so that they switched together. Uncoordinated: identical to the two-state model except that each neuron followed its own independent two-state partition (same 60 ms dwell), so switching was independent for each neuron rather than coordinated. Averaging: every neuron fired at the fixed constant mean of its two templates, 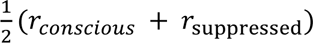, with no switching; all fluctuations in activity arose from Poisson noise. Conscious-only: every neuron fired from the conscious template, with no switching. The recorded data and all four models were then passed through the identical HMM state-inference, reconstruction, cross-neuron-validated reconstruction and reversal, and likelihood analyses, and each model was scored on whether it reproduced four signatures of the data, according to the criteria stated below. The criteria for passing each test were defined as follows. Reconstruction: in state 1, the explained variance of the conscious face significantly exceeds both zero and that of the suppressed face; in state 2, the explained variance of the suppressed face significantly exceeds both zero and that of the conscious face from the full population. Coordinated reconstruction: reconstruction survived cross-neuron validation where states were inferred from one random half of the neurons and features were predicted from the other half. Tuning reversal: the suppressed-state modulation indices were significantly negatively correlated with unambiguous modulation. As for coordinated reconstruction, states were inferred from one random half and modulation indices computed for the other half. Likelihood: in per-neuron held-out log-likelihood, a two-rate description, where each bin emitted from either the conscious or the suppressed template, exceeded the best single static rate, defined as the optimal fixed weighted average *p* × *r_conscious_* + (1 − *p*) × *r_suppressed_* of the two templates (*p* chosen by grid search of 41 uniformly spaced values from 0 to 1).

## Data availability

Electrophysiological data and associated metadata of this study will be made publicly available through the DANDI Archive upon publication.

## Code availability

Custom code used for data processing, analysis and generation of the figures will be made publicly available on GitHub upon publication.

## Acknowledgements

We thank Mo Mao, Chang-In Sohn and Yaozu Li for technical assistance, Daniel Wagenaar for help with the Neuropixels insertion design, and members of the Tsao lab for discussions.

## Author contributions

J.K.H. and D.Y.T. conceived the project and designed the study. J.K.H., F.F.L., and Y.S. collected the data. J.K.H., C.M., T.A.E. and D.Y.T. designed the analysis methods, analysed the data, and interpreted the results. J.K.H. prepared the figures. J.K.H. and D.Y.T. wrote the initial draft, which was subsequently developed into the final manuscript by J.K.H., C.M., T.A.E. and D.Y.T.

